# Live visualization of endoplasmic reticulum redox potential in zebrafish embryos reveals region-specific heterogeneity

**DOI:** 10.1101/2022.02.12.480199

**Authors:** Monika Verma, Niraj Rajesh Bhatt, Aseem Chaphalkar, Kriti Verma, Shreyansh Umale, Shweta Verma, Chetana Sachidanandan, Kausik Chakraborty

## Abstract

Redox homeostasis is an integral part of many cellular processes, and its perturbation is associated with conditions such as diabetes, aging, and neurodegenerative disorders. Redox homeostasis or redox potential in organelles is maintained within a particular range to facilitate the organelle specific cellular redox reactions. Previous studies using yeast, cell systems, and nematodes have demonstrated that the Endoplasmic Reticulum (ER) has a more oxidizing environment while the cytosol exhibits a reducing redox potential. However, we know very little about how universal this phenomenon is. We created transgenic zebrafish (*Danio rerio)* lines with roGFP sensors targeted to the ER and cytosol for studying physiological redox potential at the systems level. In the process, we also characterized the ER-targeting signal sequence in *D. rerio* for the first time. Measurements of the redox state in live embryos found that the endoplasmic reticulum exhibits deviations from its expected oxidizing redox state in different regions of the developing embryos. The ER is far more reducing than expected in certain tissues of the embryo. Cytosol also exhibited unexpected redox states in some parts of the embryo. Notably, the brain showed regions with unexpected redox states in both the ER and the cytosol. Tissue-specific differences in ER-redox potential became even more evident in a transgenic line expressing the more sensitive roGFPiE variant. Thus, live monitoring of redox potential across the developing zebrafish embryos revealed unanticipated redox states of the ER that will require new biological definitions.

## Introduction

The endoplasmic reticulum (ER) is the first stop for the proteins going through the secretory pathway and it acts as a hub for oxidative protein folding in the cell (Ellgaard & Helenius, 2003). Oxidative protein folding in the ER is mediated by redox buffers and oxidoreductases such as the Protein Disulfide Isomerases (PDIs) and ER Oxidoreductin 1 (ERO1), aided by the oxidizing redox potential in the ER (Ellgaard & Ruddock, 2005). While the ER is oxidizing, other compartments like the cytosol and the mitochondrial matrix maintain a reducing environment to prevent the resident enzymes from oxidation and to facilitate metabolic reactions (Gutscher et al., 2008; Hanson et al., 2004; Hwang, Sinskey, & Lodish, 1992; Østergaard, Tachibana, & Winther, 2004). Differences in the redox potential of the ER and cytosol can also be attributed to differences in ratios of reduced and oxidized glutathione (GSH:GSSG). The cytosol has many fold higher concentrations of GSH:GSSG (10:1 to 100:1) as compared to the ER (3:1 or 1:1) (Hwang, et al., 1992; Montero, Tachibana, Rahr Winther, & Appenzeller-Herzog, 2013; Morgan et al., 2013). However, maintaining redox potential is a complex process and apart from enzymes it also depends on the redox buffers present in the cell. The generation of live probes for redox sensing such as the roGFP variants have revolutionized the field. These sensors, targeted to various cellular organelles have been used to study intracellular differences in redox potential in a number of model organisms such as *Saccharomyces cerevisiae, Caenorhabditis elegans*, and mammalian cell lines (Ayer et al., 2013; Birk et al., 2013; Hu, Dong, & Outten, 2008; Romero-Aristizabal, Marks, Fontana, & Apfeld, 2014).

The ER is multi-functional; in addition to protein folding, it also serves as a compartment for calcium storage and lipid biosynthesis (Glaumann, Bergstrand, & Ericsson, 1975; Inui, Saito, & Fleischer, 1987; Lai, Erickson, Rousseau, Liu, & Meissner, 1988; Vidugiriene & Menon, 1993). Within a cell, the ER maintains distinct sub-regions for these functions (Baumann & Walz, 2001; Sitia & Meldolesi, 1992). Additionally, the ER in different organs in the body have different demands on it (Zheng & Staehelin, 2001) e.g. the pancreatic cells are secretory in nature with nearly 1/3^rd^-1/5^th^ of the proteome passing through the ER, making protein folding the major activity (Ghaemmaghami et al., 2003). Whereas, the ER in muscles serves more as a calcium storage hub (Burdakov, Petersen, & Verkhratsky, 2005; Coussin, Macrez, Morel, & Mironneau, 2000; Takeshima, 2002). Given this, we speculated that the ER in different tissues would adapt to the differences in demand and would exhibit differences in redox potential.

Studies in the fruit fly larvae (*Drosophila melanogaster*) and nematodes (*C. elegans*) have shown differences in cytosolic redox potential across different regions of the animal (Albrecht, Barata, Grosshans, Teleman, & Dick, 2011). Apfeld and colleagues showed that insulin signaling was important for the regulation of redox potential in the cytosol of *C. elegans* (Romero-Aristizabal, et al., 2014). Morimoto and group demonstrated that the ER becomes less oxidizing while the cytosol becomes less reducing in aging worms (Kirstein et al., 2015). Most of these studies on redox potential, performed in invertebrate models, have focused on perturbations due to chemical or environmental cues, with the assumption that the ER redox potential is constant across tissue types (Kirstein, et al., 2015). Zebrafish roGFP and HyPer7 reporters have been used to study redox events in the mitochondria, cytosol and nucleus during axonal injury (O’Donnell, Vargas, & Sagasti, 2013), fin wounding (Pak et al., 2020) and in cardiomyocytes (Panieri, Millia, & Santoro, 2017). However, we know very little about the physiological redox states of endoplasmic reticulum in the different cell types in a zebrafish embryo.

We chose zebrafish (*Danio rerio)* as the vertebrate model to explore the redox potential dynamics of the ER and cytosol, because the transparency of the embryos allows us to monitor redox states in the live animal using sensors such as roGFP. The embryonic anatomy and molecular pathways are well conserved from zebrafish to human. The chemical permeability of the embryos and larvae also make it easy to expose them to small molecule perturbation. In this study, we generated transgenic zebrafish lines ubiquitously expressing the redox sensor, roGFP, targeted to the endoplasmic reticulum (ER) and cytosol. During this process, we also characterized the ER-targeting signal sequence in zebrafish for the first time. Overall, the ER and cytosol in most tissues had the expected redox states. However, there were interesting deviations from the norm in the brain and some other tissues. These differences in redox states became even more evident when we used roGFPiE, a more sensitive variant of roGFP. Thus, our organelle-targeted redox sensors allowed us to visualize and quantify the redox states of cytosol and ER in various tissues in a live vertebrate model. These tools we have generated would be valuable in exploring the mechanisms for maintaining different redox states in different tissues and would also allow us to understand the changes in redox states induced under pathophysiological conditions.

## Results

### Generation and characterization of cytosol and ER targeted roGFP transgenic lines

To investigate if redox potential of the cytosol and the ER varies across different tissues of zebrafish, we generated zebrafish transgenic lines expressing roGFP2 either in the cytosol or the ER (Fig. 1A). The roGFP2 sensors have disulfide bonds engineered in the proteins such that the protein maintains different conformations in reducing and oxidizing milieu. Both the conformations absorb energy at 405 and 488 nm of light and emit at 505 nm, however the ratio of absorbance at these two wavelengths shifts according to the redox environment in which it is present. Absorbance at 405 is higher when the probe is oxidized (therefore the emission at 505 is higher when excited at 405 nm), while absorbance at 488 nm increases when it is in reduced state (Hanson, et al., 2004). So, the values obtained for emission at 505 nm when excited at 488 nm divided by the emission at 505 nm when excited at 405nm (488 nm/405 nm) acts as a surrogate for the redox status. Fusing roGFP2 to proteins like Grx1 is known to increase the time resolution of recording changes in redox potential (Bhaskar et al., 2014) (Gutscher, et al., 2008). These fused probes equilibrate faster with GSH:GSSH dependent redox potential and are beneficial (and sometimes necessary) for time kinetics experiments. However, we did not want to measure kinetics in this particular study, so we used native roGFP probes.

**Figure 1:**
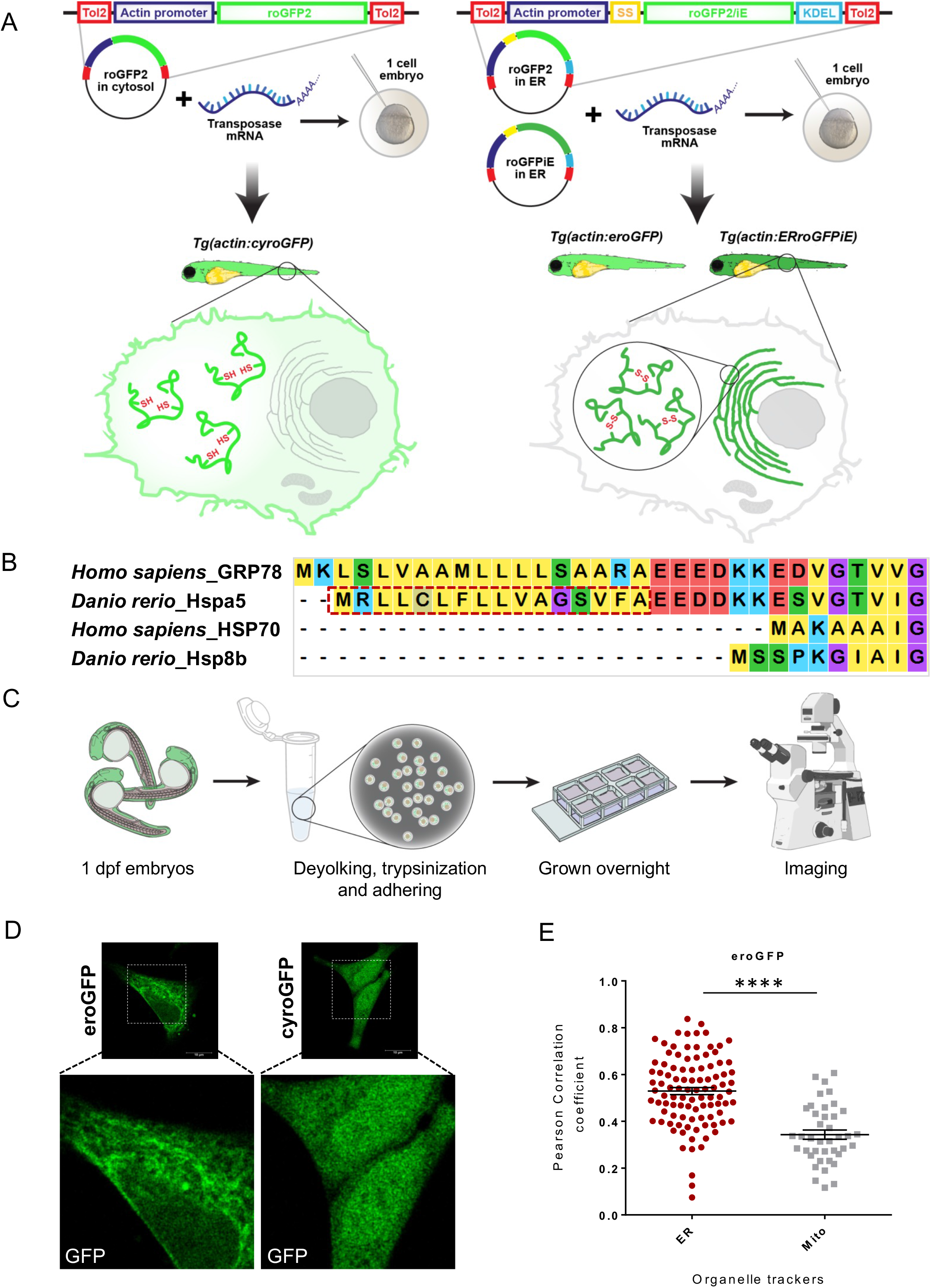
roGFP2 is localized in Endoplasmic reticulum in the eroGFP transgenic line. (A) A schematic representation of the construct and strategy used to make the transgenic lines, where roGFP2 was targeted to either cytosol or ER, and roGFP1/iE to ER. The construct consisted of actin promoter driving the roGFP2/roGFP1/iE flanked by tol2 sites on either side for correct transposase-mediated recombination. The lines are named according to the zebrafish convention of nomenclature. (B) Amino acid (aa) sequence alignment of Hspa5 (GRP78) and Hsp8b (HSP70) from zebrafish and human. Alignment was done using MEGA to find the ER targeting signal sequence specific to Hspa5 and not present in the cytosolic Hsp8b. Dotted red box shows the 16 aa used in this study for targeting roGFP2 to the ER. For ERroGFPiE line, 26 aa were taken. (C) A schematic depicting the procedure of generating primary cell culture from zebrafish embryos. Briefly, the single cell suspension from entire embryos was grown in L15 media at 28°C in chambered cover glasses followed by imaging next day. (D) Fluorescence images of primary cells made from 1 dpf zebrafish embryos showing roGFP signal in eroGFP (left panel) and cyroGFP (right panel) transgenic lines. Images in the bottom panel show enlarged areas of the cells in the top panel (confocal microscope, 63X objective, 4X zoom, scale 10µm). (E) Scatter plot shows Pearson correlation coefficient for colocalization of green and red channel for eroGFP line, where each dot corresponds to one ROI. Significance was tested using two-tailed Student’s t test with unequal variance (ROI-region of interest, ER-endoplasmic reticulum, mito-mitochondria, **** is p value <0.0001). (dpf-days post fertilization, ER-endoplasmic reticulum, mito-mitochondria, ROI region of interest)

roGFPiE is a redox sensor engineered to sense relatively oxidizing compartments like the ER (Lohman & Remington, 2008), so we made another transgenic line targeting roGFPiE to the ER. We generated three different zebrafish transgenic lines expressing roGFP2 in the cytosol, *Tg(actin:cyroGFP)*] (hereafter cyroGFP), and ER, *Tg(actin:eroGFP)* (hereafter eroGFP), and roGFPiE in the ER, *Tg(actin:ERroGFPiE)* (hereafter ERroGFPiE). The actin promoter ensured ubiquitous expression of GFP in the developing embryos (Fig. S1A). The default localization of the roGFP2 expressed would be in the cytosol. However, to target the sensor to the ER we needed to fuse an ER targeting signal sequence to the N-terminus of roGFP2. No previous study has targeted exogenous proteins to the ER in the zebrafish and thus the ER targeting signal for zebrafish was not known. We chose to identify the sequence from GRP78 or Bip, a canonical ER chaperone whose targeting signal sequence is routinely used to target proteins to the ER in *S. cerevisiae* and mammalian cells (Kanekura, Ishigaki, Merksamer, Papa, & Urano, 2013; Merksamer, Trusina, & Papa, 2008). We aligned the protein sequences of GRP78 and cytosolic chaperone HSP70 from *Homo sapiens* to that of the *D. rerio* Hspa5 and Hsp8b (HSP70) to identify the putative ER signal sequence. We selected the first 16 amino acids (aa) that appeared to be specific to the ER-targeted Hspa5 and was not present in its cytosolic counterpart, Hsp8b (Figure 1B). These 16 aa of Hspa5 were fused to the N-terminal of roGFP2 and the first 26 aa of Hspa5 were fused to the N-terminal of ERroGFPiE to assess the minimal sequence needed for targeting (Figure 1B). In order to look for the efficiency and cleavage site prediction, we used SignalP 6.0 tool for both of the signal peptides (Teufel et al., 2022). We found that signal peptide cleavage site prediction was after 16^th^ a.a. in both the cases (Figure S2A and B). Thus, the design principles of the signal sequence were, in principal, sufficient to target the protein to the ER.

In the embryos, where cells are packed together, it was difficult to visualize the organelles clearly. Thus, we prepared a primary cell culture from 1 dpf (days post fertilization) zebrafish embryos by dissociating the cells from the embryo and allowing them to attach to a substratum (Figure 1C). The GFP fluorescence in the cyroGFP cells was present throughout the cell reminiscent of localization in the cytosol (Figure 1D and S3A). The GFP fluorescence in the eroGFP showed a tubular pattern distinct from the cyroGFP; we observed a perinuclear distribution of the GFP with nuclear exclusion (Figure 1D and S3A). We co-stained the cyroGFP, eroGFP and ERroGFPiE cells with various organelle trackers to visualize organelles clearly. Visual analysis showed a similar pattern of roGFP and ER-Tracker Red in the eroGFP line; the MitoTracker Red and Lyso-Tracker Red had very different staining patterns to roGFP (Figure S3A) (Figure S4). This was confirmed by quantification. Pearson correlation coefficient analysis for colocalization of roGFP with ER and MitoTracker Red in eroGFP cells also revealed significant correlation with ER Tracker Red (Figure 1E). Similar trends were observed for the ERroGFPiE line, indicating ER localization (Figure S3C). Thus, it appears that both 16 aa as well as 26 aa can target proteins to the ER and that the 16 aa ER-targeting signal sequence is sufficient. Further, eroGFP cells also showed oxidizing environment when checked for redox status, again confirming the ER localization of roGFP (Figure 2B-D). Quantification of roGFP in the cyroGFP line did not show any colocalization with ER or MitoTracker Red (Figure S3B). We further assessed the ER localization of eroGTm was expected to cause ER stress and FP by subcellular fractionation and found significant enrichment of GFP in the ER fraction compared to nuclear fraction (Figure S3D). Due to similar densities of ER and mitochondria, it was difficult to separate these two fractions completely using biochemical methods leading to the detection of roGFP in the ER-contaminated mitochondrial fraction as well. However, the biochemical and imaging data, taken together, indicated that the redox sensitive GFPs were localized correctly in their respective compartments.

**Figure 2:**
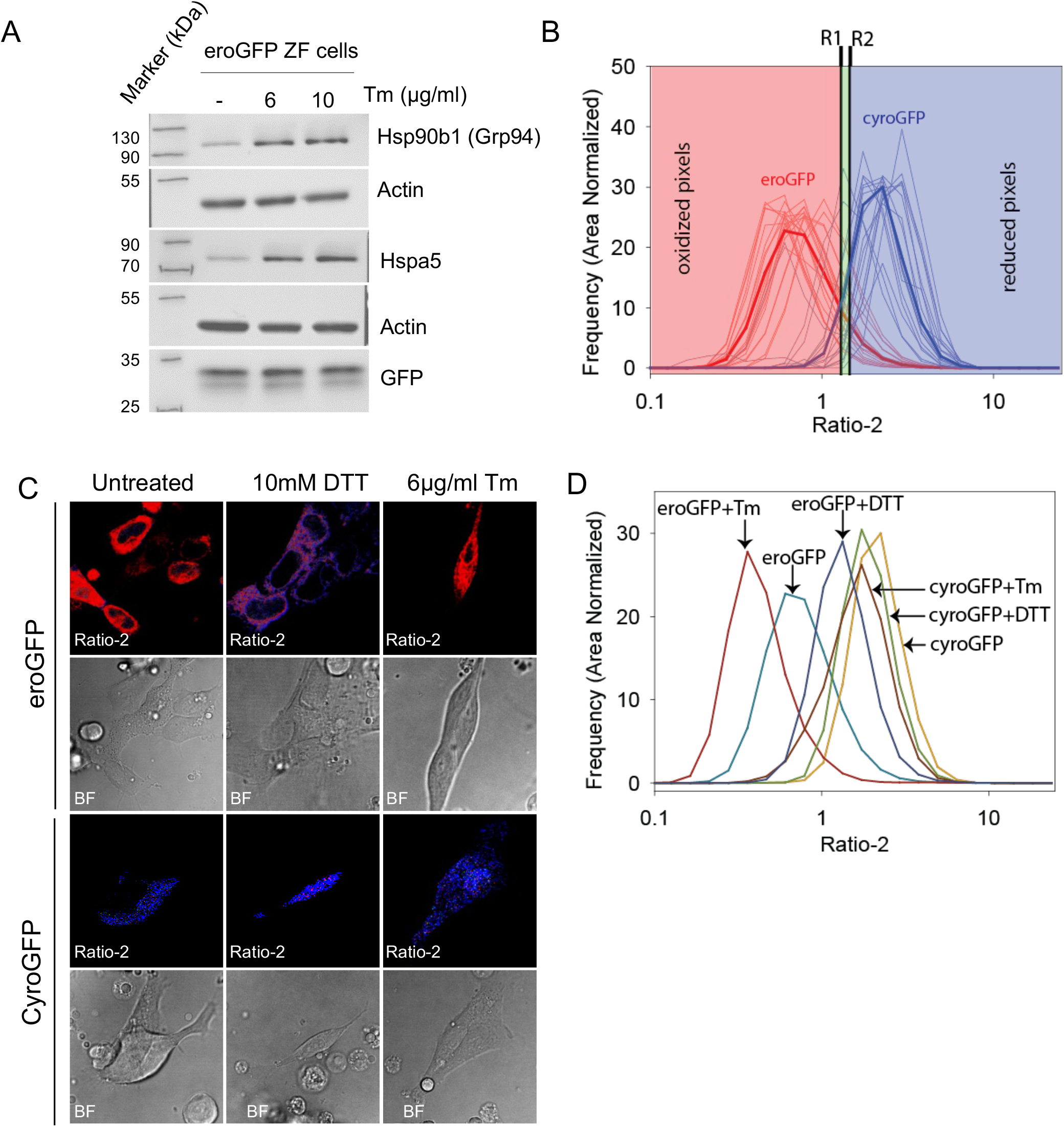
Primary cells from roGFP transgenic lines are sensitive to chemical perturbations. Primary cells from 1 dpf embryos were grown overnight and 6 and 10 μg/ml of tunicamycin (Tm) treatment was given for 8 hrs. (A) Western blots show expression of GRP78 (Hspa5) and GRP94 (Hsp90b1) as ER stress markers, actin was used as a loading control and GFP was probed to show expression of the transgenic protein. (B) Both untreated and treated cells were imaged to determine redox potential using confocal microscope. Raw images were then analyzed by MATLAB script to calculate pixel wise Ratio-2 values. Frequency of Ratio-2 values was area normalized for untreated cells and graph was plotted. The pixels were divided into 3 bins according to the values obtained and given colors in RGB scale (Red-Green-Blue). R1 is the boundary defined as 1 standard deviation away from mean eroGFP value while R2 is 1 standard deviation less than mean cyroGFP value. Ratio-2 values below R1 were marked red (oxidizing), between R1 and R2 marked green (intermediate) and higher than R2 were marked blue color (reducing). (C) Cells were treated with either 6 µg/ml of Tm (8 hrs of treatment) or 10 mM DTT (immediately imaged after treatment). Images represent maximum projection images of untreated, 10 mM DTT and 6 µg/ml Tm treated cells from left to right respectively (Confocal microscope, 63X objective, zoom 4, scale 10 µm). (D) Ratio-2 values are plotted for each of these conditions. Each line represents mean Ratio-2 for each treatment (SET1), done on one day and it was repeated thrice. N is variable in each condition and contains value from at least 6 fields. (BF-bright field)

### The eroGFP line does not exhibit ER stress

Overexpression of exogenous proteins can perturb and stress the cell. We assessed the eroGFP lines for ER stress marker expression. The expression of GRP78 (Hspa5 in zebrafish), known to be upregulated during ER stress (Kozutsumi, Segal, Normington, Gething, & Sambrook, 1988), did not change significantly in the eroGFP line compared to WT (Figure S5A and B). Manganese superoxide dismutase (SOD2) protects cells against oxidative stress (Kasahara, Lin, Ho, & Reddy, 2005; Wang, Branicky, Noë, & Hekimi, 2018). Our studies showed no significant change in Sod2 expression in the transgenic lines (Figure S5A and C). An increase in disulfide bond formation in the ER due to overexpression of proteins may also result in increase in the reactive oxygen species (ROS) levels (Harding et al., 2003; Zeeshan, Lee, Kim, & Chae, 2016), especially since eroGFP harbors engineered disulfide bonds (Hanson, et al., 2004). We used the DCFDA assay to detect ROS levels and found it to be comparable between the eroGFP and the control embryos (Figure S5D). Thus, we concluded that the eroGFP transgenic lines did not show any signs of ER stress in normal conditions.

### The roGFP lines respond to external perturbations in cellular homeostasis

For the roGFP transgenic lines to be useful for monitoring redox potential in the animal, the roGFP proteins must be responsive to changes in the oxidative state. We measured the emission peak of the roGFP when excited at 405 nm and 488 nm with live emission scans of 4 dpf larvae on the confocal microscope. We found the expected 505 nm peak when excited at 405 nm or 488 nm in all transgenic lines (Figure S6A-E). The head region in eroGFP embryos showed peak at 505 nm upon excitation at 405 nm or 488 nm indicating that roGFP is oxidized in the ER, while cyroGFP embryos showed very minimal 405 signal (505 nm emission with 405 nm excitation) suggesting a reducing milieu (Figure S6A and C). Similar results were also obtained for the trunk region of the larvae (Figure S6B and D). Thus, roGFP2 targeted to ER and cytosol reported the expected relative redox differences between the two compartments. The transgenic line ERroGFPiE also showed two characteristic peaks where the emission signal at 405 nm was higher than 488 nm, as previously reported for this variant (Figure S6E) (Lohman & Remington, 2008). The oxidized form of roGFPiE is predominant in the ER (405 peak in Figure S6E). These peaks were absent in the WT zebrafish embryos (Figure S6F) showing the signals to be specific for the fluorescent reporters.

To assess the response of the roGFP lines to perturbations in cellular redox potential we used dithiothreitol (DTT) and tunicamycin (Tm) treatment. To avoid variability due to drug penetration in the embryo, we prepared a primary culture of cells dissociated from 1 dpf embryos. The cells were treated either with 10 mM DTT (reducing reagent) or 6 µg/ml and 10 µg/ml of Tm (ER stress inducer). To test the efficacy of Tm in inducing ER stress in the primary culture cells we collected protein from the cells after 8 hours (hrs) of treatment and performed western blots for ER stress markers. We found upregulation of both Hspa5 and Hsp90b1 (GRP94) (canonical ER stress markers) in the eroGFP cells at both the concentrations of Tm (Figure 2A). All further perturbation experiments were performed after 8 hrs of treatment with 6 µg/ml of Tm. All DTT treatments were done with 10 mM DTT, and imaging was started within 1-2 minutes (min) of addition of the compound.

Two different fluorescence measurements were done for both the eroGFP and cyroGFP probes: fluorescence emission from 505-540 nm upon excitation at 488 nm and from 505-540 nm upon excitation at 405 nm. The fluorescence ratio of these two measurements 488 nm/405 nm were then obtained after analysis and will be henceforth referred to as Ratio-2. Pixel wise frequency distribution of Ratio-2 in eroGFP and cyroGFP cells showed that they followed a roughly log-normal distribution (Figure 2B). We transformed the ratio in log2 scale and set two boundaries for quantitative demarcation of reduced and oxidized regions. R1 boundary was defined as 1 standard deviation more than the mean eroGFP value while R2 was set at 1 standard deviation less than mean cyroGFP value. Pixels with Ratio-2 less than R1 represented ‘oxidized’ pixels and those with Ratio-2 greater than R2 represented ‘reduced’ pixels. Pixels between R1 and R2 represented the intermediate between oxidized and reduced state (Figure 2B). For ease of visualization in the cellular and embryonic images, the oxidized pixels were false-colored red, the reduced pixels were colored blue and the intermediate pixels green.

We determined the Ratio-2 in the transgenic cells treated with DTT. Treatment of eroGFP cells with DTT resulted in instant and significant increase in the Ratio-2, indicating a change from oxidized to reduced state (Figure 2C and D). Tm was expected to cause ER stress and induce reducing conditions in the ER (Merksamer, et al., 2008). Later, the work by same group reported reflux of roGFP to cytosol upon Tm treatment, being the reason for observed reducing signal (Igbaria et al., 2019). Instead, treatment with Tm decreased the Ratio-2 in eroGFP marginally suggesting a shift to oxidized state in our study (Figure 2C, 2D and S7). This trend towards a decrease in the Ratio-2 when treated with Tm was not consistent across different sets of cells (Figure S7); importantly the ER redox potential did not show a shift towards reducing state upon treatment with Tm. Importantly there was no redistribution of eroGFP from ER to the cytosol. This data also suggests that reflux of roGFP is not prominent in our model. The cyroGFP cells did not show any significant change in the Ratio-2 in response to DTT or Tm; likely because the cytosol is already reduced (Figure 2C, 2D and S7). Taken together, the eroGFP and cyroGFP zebrafish lines were able to report the expected redox potentials and respond to external perturbations.

### Redox potential of ER and cytosol show region wise differences

Obtaining a quantitative redox map of zebrafish embryos was fraught with hurdles. High autofluorescence (scatter) in certain regions of the embryo precluded determination of the correct ratio of GFP fluorescence upon excitation at the two different wavelengths. To solve this problem, we needed to determine the contribution of autofluorescence in the different regions of the embryo. We took fluorescence images and collected emission from 505 nm – 540 nm upon excitation at either 405 nm or 488 nm using WT zebrafish embryos with no roGFP2 expression. While excitation at 488 nm had negligible autofluorescence, 405 nm showed high autofluorescence in certain regions of the embryo. Since each embryo has unique variations in anatomical structures it is impossible to create a universal quantitative map of autofluorescence in these embryos. To obtain the contribution of autofluorescence in the 405 nm excitation channel in the roGFP transgenic embryos we needed to generate an autofluorescence map for each embryo. Deconvoluting autofluorescence for a fish with a roGFP2 probe would require us to determine the contribution of autofluorescence with a proxy measurement at a wavelength where the authentic roGFP2 fluorescence has negligible contribution. We found that WT embryos when illuminated at 405 nm, showed an autofluorescence emission peak around 515 nm and exhibited a prominent shoulder between 440 nm and 480 nm (Figure 3A, upper panel). This shoulder was absent in the fluorescence peak of purified roGFP2 (Figure 3A, lower panel). Thus, autofluorescent regions in a roGFP embryo show significant emission (in the shoulder) between 440 nm and 480 nm while excited at 405 nm. This was used to differentiate from true roGFP fluorescence. We therefore recorded an additional image of each embryo with 405 nm excitation and measured the emission from 440 nm-480 nm (Figure 3B). This image allowed us to build a quantitative estimate of the autofluorescent regions of the embryo; these regions were excluded from all further analysis.

**Figure 3:**
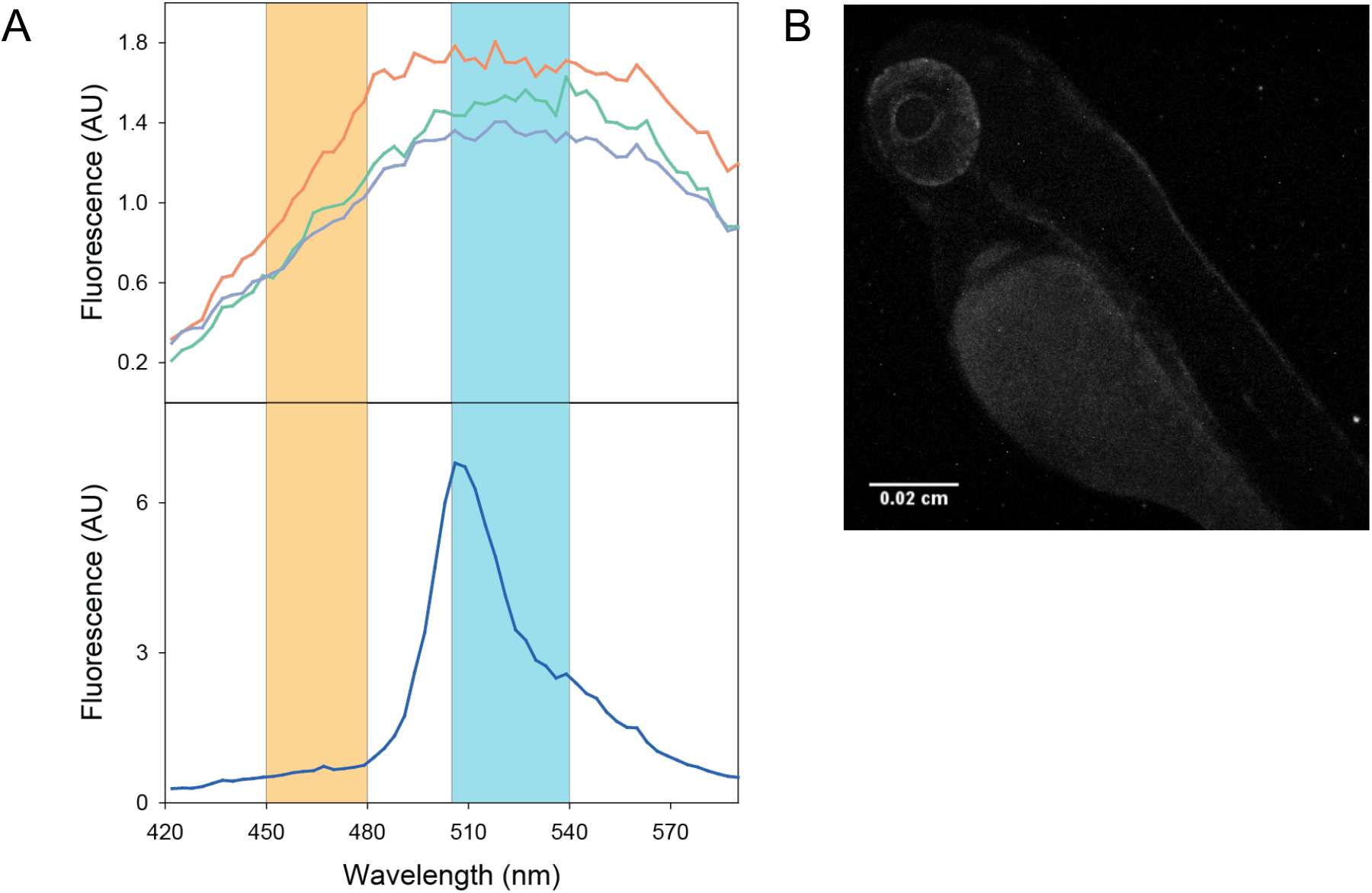
Determining autofluorescence signals from a WT embryo. (A) WT embryo lacking any roGFP expression was anaesthetized, mounted in lateral position, excited at 405 nm and emission scan was taken from 422 to 593 nm to determine autofluorescence signals (upper panel). Readings were taken at every 3 nm interval and the graph was plotted. Three lines represent three individual embryos. Similarly, emission scan was taken for 1 µM roGFP2 solution at same settings (lower panel). (B) Embryos were anaesthetized, mounted in agar in lateral position and imaging was done with excitation at 405 nm and emission was collected from 440-480 nm. Representative autofluorescence signal from one WT embryo is a maximum projection image adjusted with ImageJ. (Confocal microscope, 10X objective, scale-0.02 cm)

Another problem we faced while imaging zebrafish larvae was visual access to the whole embryo under the confocal microscope. Due to the thickness of the tissue, we could capture only one of the lateral halves of the larva. As the tail region of the animal is thinner, we could image nearly the whole depth, but the head and trunk containing the visceral organs could be only partially imaged. All confocal scans were performed with the larvae oriented laterally. The 3-D stacked images could then be rotated and viewed from various angles according to requirement (Figure S8). Optical sections were then taken to look closely beneath the surface of the regions we found important. That particular section used in each image is indicated by the schematic of a plane slicing through the 3-D reconstructed image (Figure 4A).

**Figure 4:**
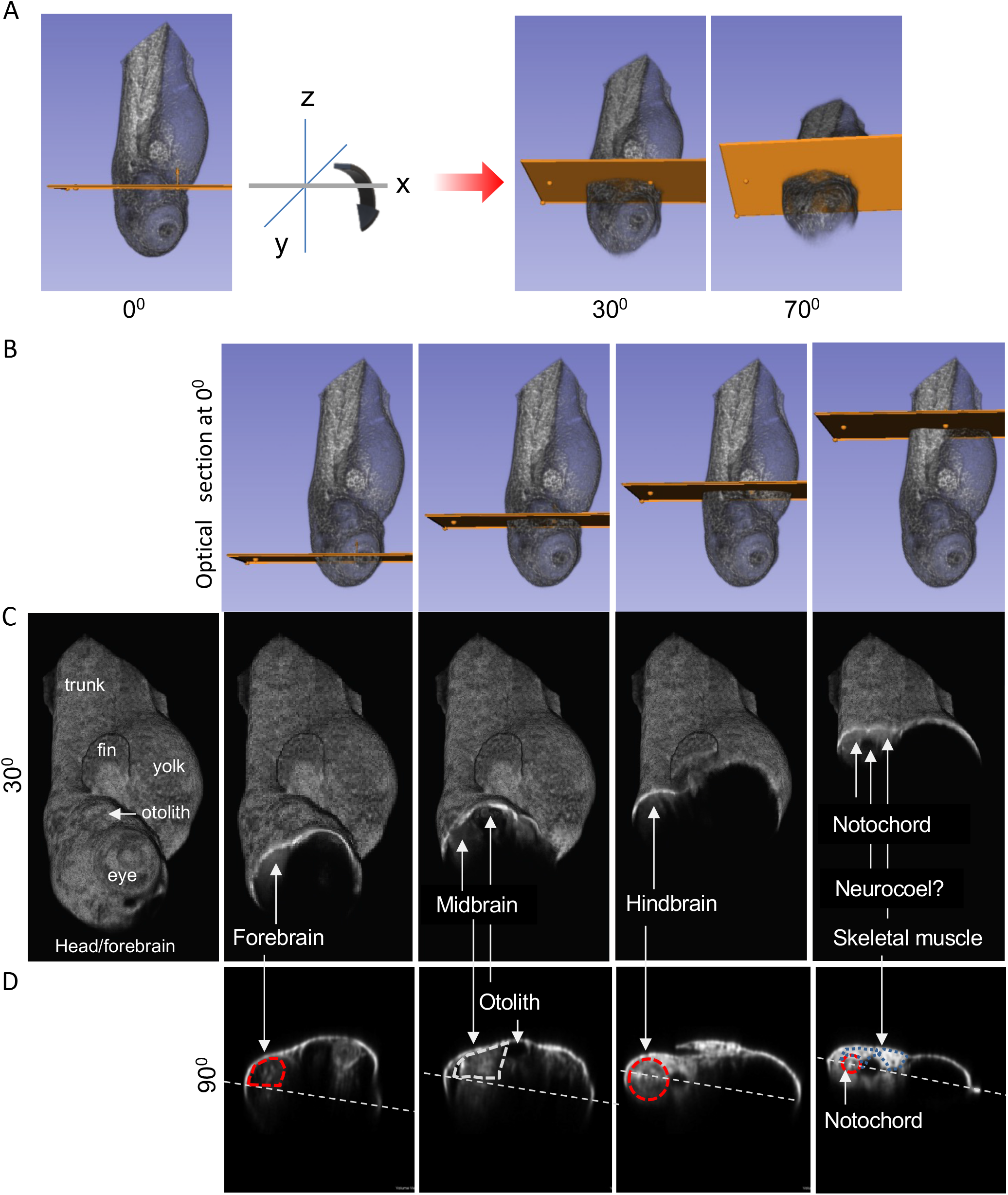
Representative cross sections and regions of interests marked on GFP fluorescence images of an embryo. (A) The left image shows the way embryo was imaged, with yellow plane showing the region where optical sectioning was done to visualize the redox ratios inside the embryo. Middle and right image indicates the way embryo was oriented (rotated) to 30^°^ and 70^°^ respectively, also indicated by the arrow, making yellow disc visible in the images. (B) to (D) Images show GFP signal of an embryo representing the rotation angles to show redox ratios in the subsequent figures. Cross section at these positions remains similar for further figures. (B) All images in the panel show the region from where optical sectioning was done. (C) All images in the panel shows the GFP signal in the oriented image of embryo sectioned at that region. (C) mages in the panel from left to right shows labeling of major regions, forebrain, midbrain, hindbrain, muscle, neurocoel and notochord, which become visible after optical sectioning done at the places indicated and embryo rotated at 30^°^. (D) Images in the panel shows same regions (as in C) when embryo is rotated at 90^°^. White line in each image indicates the lateral line which will divide the embryo in left and right halves. (Confocal microscope, 10X objective)

Once the images were acquired and analyzed, it was a challenge to identify the precise anatomical structures showing interesting redox potential patterns due to lack of reference points in the dark images. Thus, for the current study we have focused our analysis on four virtual slices (1-4) that cover the broad regions of brain and muscle in the larvae (Figure 4 B, C, and D). We have observed many interesting regions of redox anomaly outside the brain and muscle which are not discussed here. The full dataset in raw format is hosted on the Dataverse server (https://doi.org/10.7910/DVN/PRCFJM) and we hope that researchers would be able to identify their favorite regions from the dataset for further analysis.

To determine the highest and lowest boundaries of Ratio-2, we measured the Ratio-2 for a solution of purified roGFP2 in either DTT or N,N,N′,N′-tetramethylazodicarboxamide (diamide) to obtain Ratio-2 for the fully reduced form or the disulfide bonded form, respectively. While comparing the Ratio-2 distribution of the recombinant protein roGFP2 with the Ratio-2 of the eroGFP or cyroGFP embryos we observed an inconsistency. The eroGFP embryos that should ideally be fully oxidized showed a ratio lower than the fully oxidized recombinant protein (Figure S9A). Similarly, cyroGFP, ideally fully reduced, showed a ratio lower than fully reduced roGFP2 protein (Figure S9A). This suggested that the Ratio-2 distribution might be skewed to a lower value due to the residual background intensity in the 405 nm excitation channel. The fluorescence of roGFP2 in the 405 nm channel for both eroGFP and cyroGFP lines is much lower than in the 488 nm channel. This makes the denominator (during Ratio-2 calculation) more sensitive to background noise (Figure S9C and D). Importantly, the fold difference between the fully reduced and fully oxidized forms of purified roGFP2 was similar to the fold difference in Ratio-2 between cyroGFP and eroGFP in the embryos (Figure S9B). This suggested that the peaks of eroGFP and cyroGFP from the embryo must correspond to the peaks for oxidized and reduced roGFP, respectively, in the context of embryos with non-zero background fluorescence. Henceforth, instead of using the theoretical maxima and minima for Ratio-2 we decided to use the empirically determined peaks as maxima and minima for all calculations.

To be conservative in defining unexpected deviations in redox potential we used the eroGFP as controls for cyroGFP. Since the ER-targeted roGFP is considered to be in a completely oxidized state, we picked the median of the eroGFP Ratio-2 as the lowest boundary (R3) for an oxidized state of cyroGFP. Thus, we defined any pixels with a Ratio-2 less than R3 as oxidized. Pixels with Ratio-2 higher than R4 (median + one standard deviation of eroGFP) was defined as reduced cyroGFP (Figure 5A).

**Figure 5:**
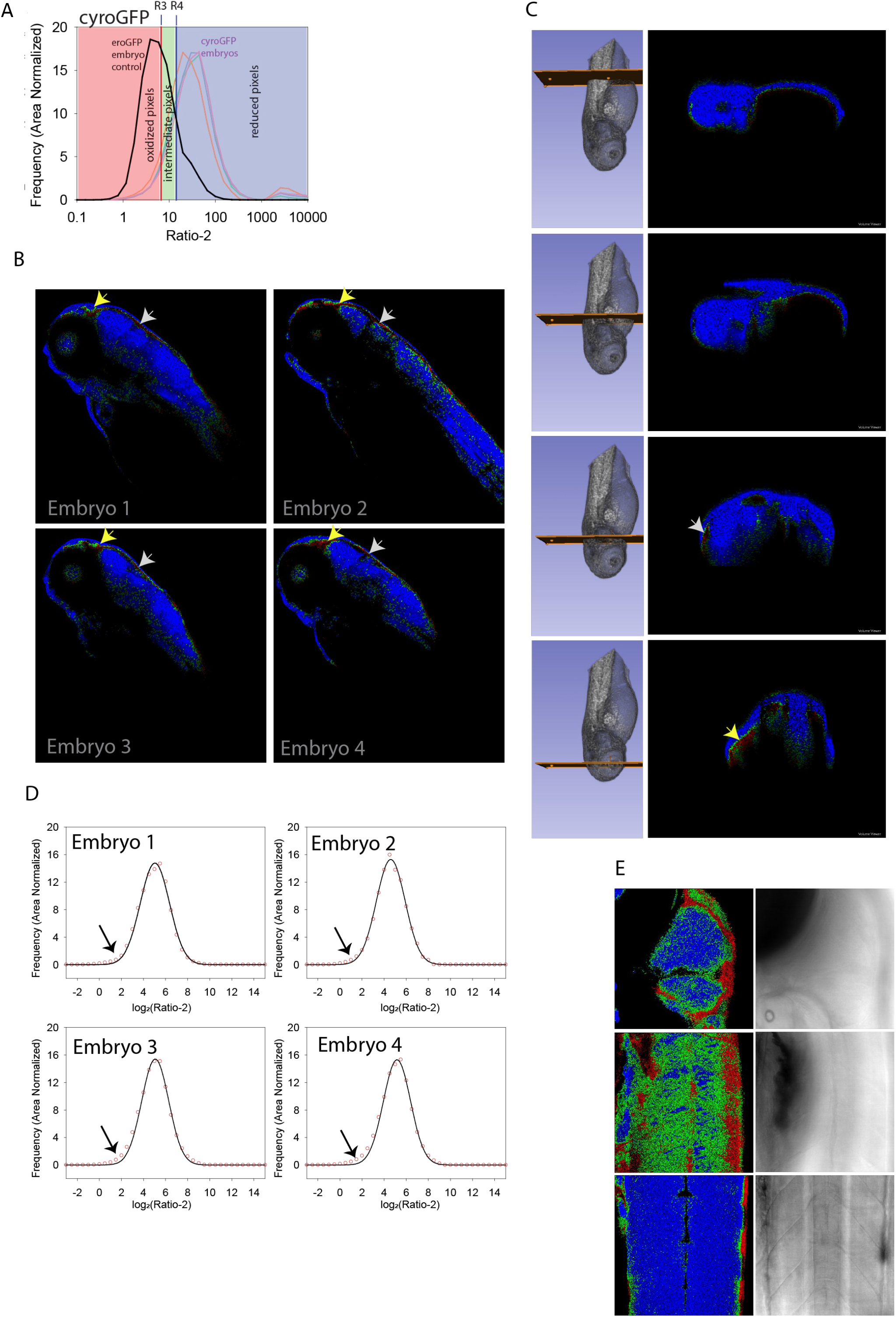
Redox status of cytosol is heterogeneous in zebrafish embryos. 3 dpf live cyroGFP embryos were anaesthetized, mounted in lateral position, and imaged by sequential scanning method, i.e., emission signal was collected from 505-540 nm first by excitation at 488 nm followed by excitation at 405 nm. Autofluorescence signal was also collected, and images were analyzed to obtain redox ratios (more details in materials and methods). (A) The frequency of Ratio-2 values for cyroGFP and eroGFP lines were plotted to show the differences in the distribution of Ratio-2. R3 and R4 boundaries were set to define color scale where R3 is median of ratio-2 from eroGFP embryo and R4 is one standard deviation from eroGFP median. Ratio-2 values below R3 were denoted as oxidizing and given red color. Ratios higher than R4 were denoted as reducing and given blue color while intermediate values between R3 and R4 were green. (B) Color coded Ratio-2 values of four embryos depicted in RGB scale as embryo 1, 2, 3, and 4. (C) Left and right panel show GFP fluorescence with region of optical section and analyzed Ratio-2 images of a representative embryo respectively. On the right panel, first, second, third and fourth panel are images obtained after optical sectioning and embryo rotation to 90°. Yellow and white arrow indicates forebrain and hindbrain respectively in (B) and (C). (D) Frequency distribution of log_2_ values of Ratio-2 were fitted with a single peak Gaussian distribution. Red colored circle represents Ratio-2 values for all embryos and black solid line is obtained after Gaussian fit of the Ratio-2 values. The black arrow shows regions where oxidizing redox ratios are above the fitted black solid line (E) Higher magnification (40X) images of a representative embryo shows the head region (behind the eye; black region on top left), initial trunk (behind the yolk) and trunk regions (muscle; myotomes) from top to bottom while left and right are Ratio-2 and BF images respectively. (BF-bright field, confocal microscope, 10X and 40X objectives)

Once we applied all the above discussed corrections in the analysis, we uncovered an interesting paradox. We found that the cytosolic redox potential was strongly oxidizing (sometimes as much as the ER) in multiple regions of the embryo (Figure 5B and C). This is supported by the frequency histogram (Figure 5D). Could these observations be spurious, or an artifact of the arbitrary division of a normal distribution? We think not, because of three reasons. One, the number of pixels in the lower range of Ratio-2 for cyroGFP was more than expected from a normal distribution obtained after fitting the curve for the top 75% of the Ratio-2 distribution (Figure 5D). Two, instead of being randomly distributed irrespective of anatomical regions, the oxidizing regions (red pixels) formed defined clusters in similar regions of the embryos, surrounded by intermediately oxidizing regions (green pixels) (Figure 5B and C). We confirmed the clustering by confocal imaging at a higher magnification (Figure 5E). Three, the oxidizing and the intermediately oxidizing regions were consistent between different embryos (Figure 5B). Thus, cyroGFP could delineate regions of the embryo with oxidizing cytosolic redox potential. Closer examination of the images (at higher magnification; 40X) revealed that the highly oxidizing regions of the embryo were absent around the skeletal muscles and were enriched in areas surrounding the midbrain and forebrain regions (Figure 5B, C, and E). This was true for multiple 3 dpf embryos (Figure 5). Close observation also revealed that the oxidizing regions formed a layer surrounding the ‘normal’ reducing regions deeper in the brain (Figure 5E). From our data it is not clear whether the oxidizing layer is within the brain tissue or is a layer outside of the brain.

### Map of the redox potential of ER using eroGFP

For analyzing eroGFP and for creating a conservative map of unexpected deviations in redox potential in the ER we used the cyroGFP peaks as controls. We defined the median Ratio-2 of cyroGFP (R6) as the boundary for identifying anomalous reduced pixels in the eroGFP embryos. Pixels with Ratio-2 less than R5 (one standard deviation lower than the median of cyroGFP) were defined as regions with eroGFP in oxidized state. It must be noted here that the boundaries (R5, R6) defined for eroGFP are distinct from the boundaries defined for cyroGFP (R3, R4) (Figure 6A). This gave us a conservative map of eroGFP redox potential.

**Figure 6:**
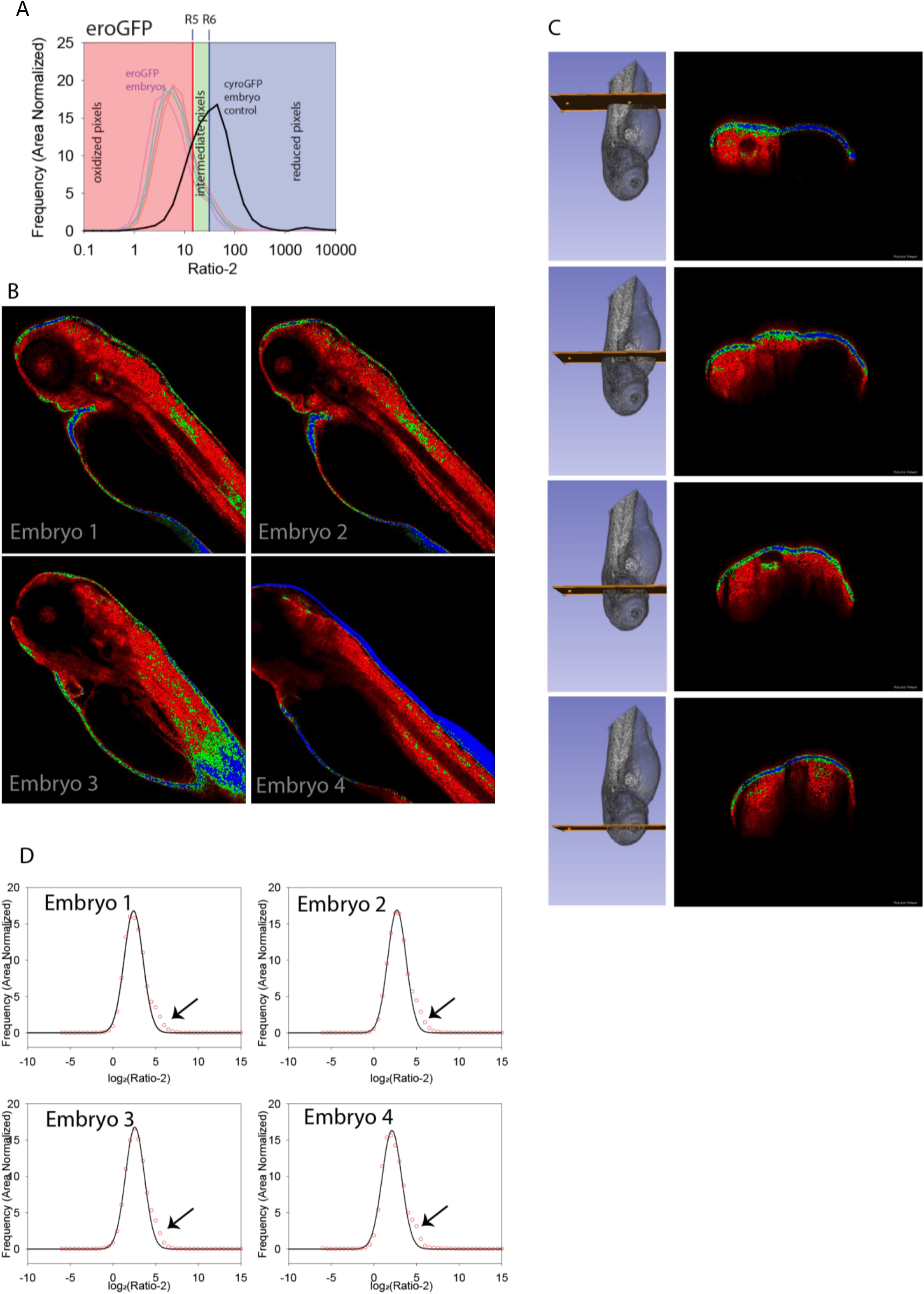
redox state of ER differs across regions of zebrafish embryos. 3 dpf live eroGFP embryos were anaesthetized and imaged to obtain redox status. Pixel wise ratios were calculated to obtain Ratio-2 values using MATLAB. (A) log_2_ of Ratio-2 values of eroGFP embryos were plotted to obtain the frequency distribution and define color scale with cyroGFP embryo as control. R5 and R6 boundaries were defined where R6 is median of cyroGFP embryo and R5 is one standard deviation lower than cyroGFP median. For color coding, pixels below R5 and higher than R6 were given red (oxidizing) and blue (reducing) color respectively while the intermediate pixels were colored green. (B) Color coded Ratio-2 images of four embryos are shown separately. (C) Left and right panel show GFP fluorescence with region of optical sectioning and analyzed Ratio-2 images respectively. First, second, third and fourth panel from top to bottom show the ratio-2 images sectioned at the indicated positions and rotated to 90°. (D) log_2_ values of ratio-2 were plotted against frequency with area normalization where red circles are the actual calculated values while black solid lines are fitted values. The graphs were made for each embryo individually. Black arrow indicates the real Ratio-2 values above the fitted data. (Confocal microscope, 10X objective)

As expected, ER in most regions of the embryo was oxidizing in nature (Figure 6B and C). However, a transverse section of the 3D reconstructed images showed a layer with ER in a strongly reducing state just below the skin of the embryo. We checked the Ratio-2 distribution of eroGFP embryos and found that indeed there were many pixels in the intermediate and strongly reducing state than expected from a normal distribution shown by fitted black line (Figure 6D). Transverse section slices confirmed that the strongly reducing pixels (blue/green) were present as a layer surrounding the embryo, suggesting a layer of cells on or below the skin (Figure 6C). Small defined regions with pixels in the intermediate reducing state were present in the hindbrain and forebrain (Figure 6B and C (last two panels)). These results suggested that there are regions in the embryo that harbor ER with lower redox potential than expected. However, roGFP2 is not a very sensitive probe for ER potential measurement because it is oxidized well below (−272 mV) the redox potential of ER (−180 mV to −235 mV). Thus, we chose the ERroGFPiE line to characterize differences in ER redox potential across different regions of the embryo.

The ERroGFPiE probe indeed accentuates the deviations-from-expected in the redox potential of the ER across the embryo as seen from the frequency distribution of Ratio-2 for ERroGFPiE (defined as Ratio-iE henceforth) (Figure 7A). Distribution of Ratio-iE did not fit well to a single peak or a double peak normal distribution and consists of at least 3 underlying normal distributions consistent among the different embryos tested (Figure 7A). The peak of the distribution of lowest Ratio-iE corresponds well with the ratio obtained from completely oxidized roGFPiE whereas the peak of the highest distribution corresponds well with the completely reduced roGFPiE (Figure 7A). Thus, roGFPiE does not suffer from the drawback of autofluorescence that shifted the ratio of eroGFP embryos towards lower values. This is primarily due to higher fluorescence of ERroGFPiE in the 405 nm channel as also reported for roGFPiE variant (Lohman & Remington, 2008). Conservation of the distributions among different embryos also reinforces the observation that there are hyper-reducing regions in the embryos.

**Figure 7:**
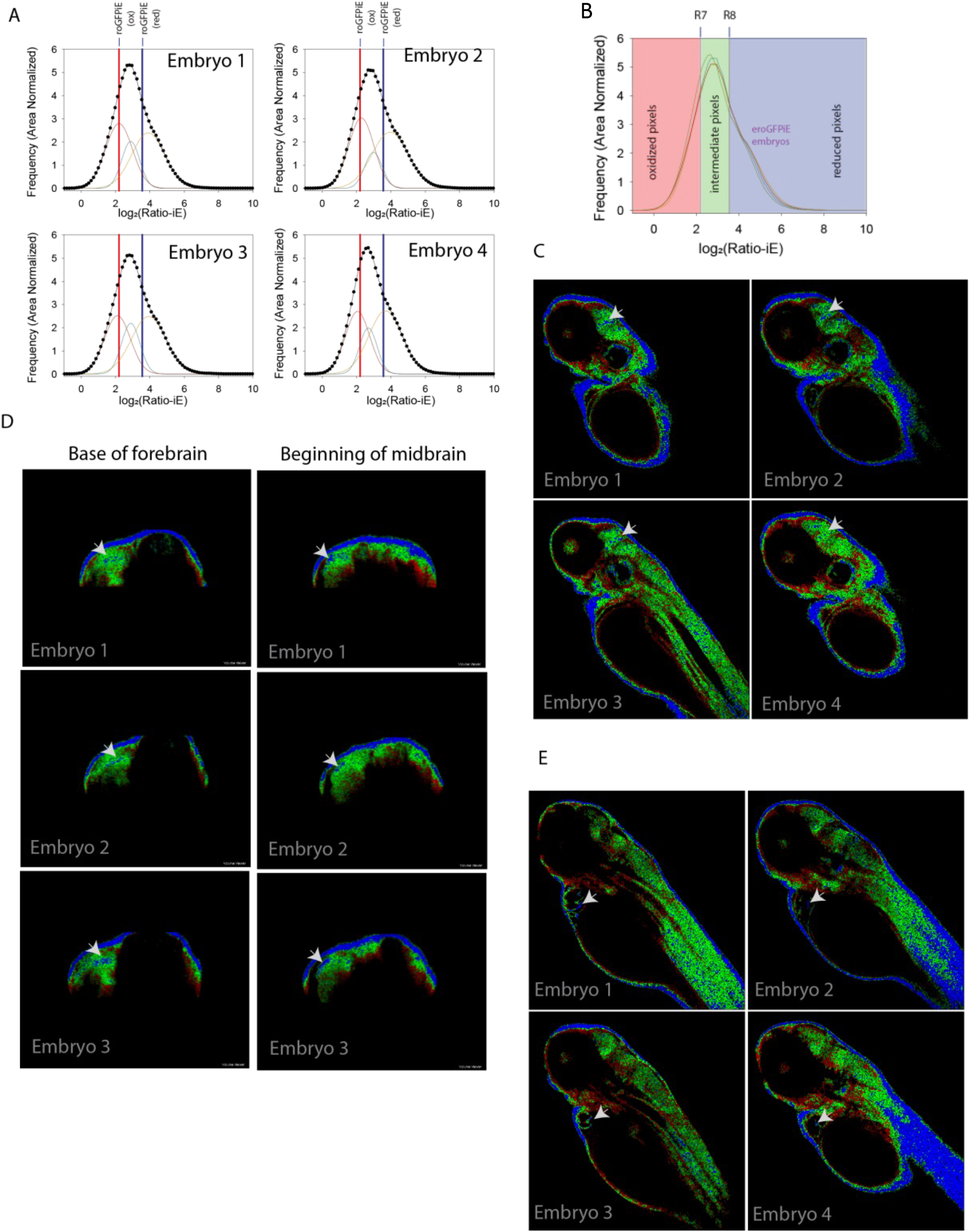
The more sensitive ERroGFPiE line reveals heterogeneity in the ER redox status across regions of the embryo. 3 dpf live ERroGFPiE embryos were anaesthetized, mounted in soft agar in lateral position, imaged, and analyzed for redox (Ratio-iE) values. The roGFPiE solutions treated with 10 mM DTT or 10 mM diamide were also imaged using same settings, i.e., emission collection from 505-540 nm when first excited at 488 nm and then at 405 nm. (A) Frequency distribution for the log_2_ values of Ratio-iE for ERroGFPiE embryos were fitted to a combination of three normal distributions. The fitted peaks are shown by three smaller peaks where red and blue line represents the Ratio-iE of fully oxidized (diamide treated) and reduced (DTT treated) roGFPiE solutions respectively. Individual graphs were plotted for the 4 embryos. (B) The log_2_ Ratio-iE distribution for ERroGFPiE embryos was binned in three groups using reduced and oxidized Ratio-iE values from roGFPiE solutions. Ratios lower than the median of oxidized roGFPiE solution (R7 boundary) were oxidizing and given red color. The values higher than the median of reduced roGFPiE solution (R8 boundary) were given blue color (reducing regions) and intermediate values between R7 and R8 were given green color. (C) Ratio-iE images of four roGFPiE embryos are shown using the described color scale where white arrow shows the reducing patches in the forebrain. (D) Ratio-iE images obtained after optical sectioning at forebrain (left panel) and midbrain (right panel) regions followed by rotation of the embryos to 0° are shown. White arrow indicates reducing patches in the base of forebrain and beginning of the midbrain in three independent embryos (top to bottom). (E) Ratio-iE images of the embryos in a different Z plane than C shows reduced status in the heart walls (white arrows). (Confocal microscope, 10X objective)

We defined the cutoffs for Ratio-iE based on the ratios obtained from measurements of the purified roGFPiE protein. Pixels having Ratio-iE more than the median Ratio-iE of reduced roGFPiE (DTT treated; R8 boundary) were marked as reducing regions (blue pixels), whereas pixels having Ratio-iE lower than that of oxidized roGFP-iE (diamide treated; R7 boundary) marked regions with oxidizing redox potential (red pixels) (Figure 7B). The mild deviations seen in the eroGFP embryos became clearer in ERroGFPiE as patches of reducing regions in the forebrain (blue pixels) (Figure 7C). From the transverse optical sections, we could see conserved regions of reducing state at the caudal limits of the forebrain and the surface of the midbrain (Figure 7D). We also observed reducing regions possibly around the pericardium (Figure 7E). Images were taken at higher magnification to confirm the locations of the reducing regions in the ERroGFPiE embryos (Figure S10A and B). At 40X magnification we were able to observe similar striations of reducing regions in the forebrains (Figure S10B). The trunk region also had striations of reducing regions paralleling the chevron-shaped muscles (Fig. S10B).

Our results suggest that the ER may not be as uniformly oxidizing throughout the embryo as previously thought (Figure 7A). The distribution of Ratio-iE in the embryos corroborates well with the heterogeneity seen in the redox potential of the ER. The lowest peak in the distribution coincides with completely reduced roGFPiE. The middle peak corresponds to a redox potential of ~ −234mV (Table 1) and is consistent between the different embryos. This peak corresponds roughly to the reported redox potential of the ER as reported in some of the studies (Delic, Mattanovich, & Gasser, 2010). Roughly 26 + 11% of the pixels represent the ER in this state, a similar percentage remains in a more oxidized (~22+12%) state but the most number of pixels represent the reduced (~51+14%) state (Table 1). Minor interindividual variations are captured in the data along with some differences in experiments performed on different days (Table 1). However, the data largely remains consistent over days and across embryos, showing the robustness of the approach. This data along with the images suggest that there is not a single universal redox potential of the ER in the embryo. It is important to note that the pseudo-coloring is conservative and isolates only high-confidence hyper-oxidizing and hyper-reducing regions. Since the middle peak significantly overlaps with the low and high peaks it is difficult to illustrate the range of heterogeneity using pseudo-colors. The heterogeneity in redox potential may explain the range of redox values of the ER that have been measured in single cell organisms or in cell lines (Avezov et al., 2013; Brach et al., 2009; Schuiki, Zhang, & Volchuk, 2012; van Lith, Tiwari, Pediani, Milligan, & Bulleid, 2011). Taken together, our study revealed that the redox state of cytosol and ER across the vertebrate embryos is not uniform and has many regions of heterogeneity.

### ER redox potential is robustly maintained when proteostasis is challenged

To check if the redox distribution is significantly perturbed when ER proteostasis or cellular proteostasis is perturbed, we treated the 2 dpf embryos with chemicals that alter ER-proteostasis (Tunicamycin, Tm) or global cellular proteostasis (L-azetidine-2-carboxylic acid, AZC) and imaged at 3 dpf. We also checked for the Hspa5 upregulation upon Tm treatment by western blotting and we found a significant increase in Hspa5 as compared to the WT control (data not shown). Similarly, we found Hsp90 upregulation upon AZC treatment validating the effectiveness of the drug (data not shown). Interestingly, AZC did not change the distribution or the mid-point redox potential of the ER (Table 2) (Figure 8A). By contrast, alteration of ER-proteostasis by Tm increased embryo to embryo variability of redox status (as seen from the large standard deviation) (Table 2) (Figure 8B). The treatment also altered the distribution with significant increase in percentage of voxels in oxidized (OX) bin when compared to untreated control embryos imaged on the same day. There was a significant decrease in reducing (RED) voxels too; however, we observed high biological variability as well (Table 2) (Figure 8B). The mid-point redox potential exhibited a slight decrease and shifted towards reducing potential, but this change was not significant either (Table 2). This indicates that the ER redox potential is robust and is maintained at constant values during challenges to proteostasis. However, the inter-individual differences are more during ER-stress indicating that some of the embryos were more affected by Tm induced ER stress than others. It will be important to understand the underlying basis of these interindividual differences during ER stress.

**Figure 8:**
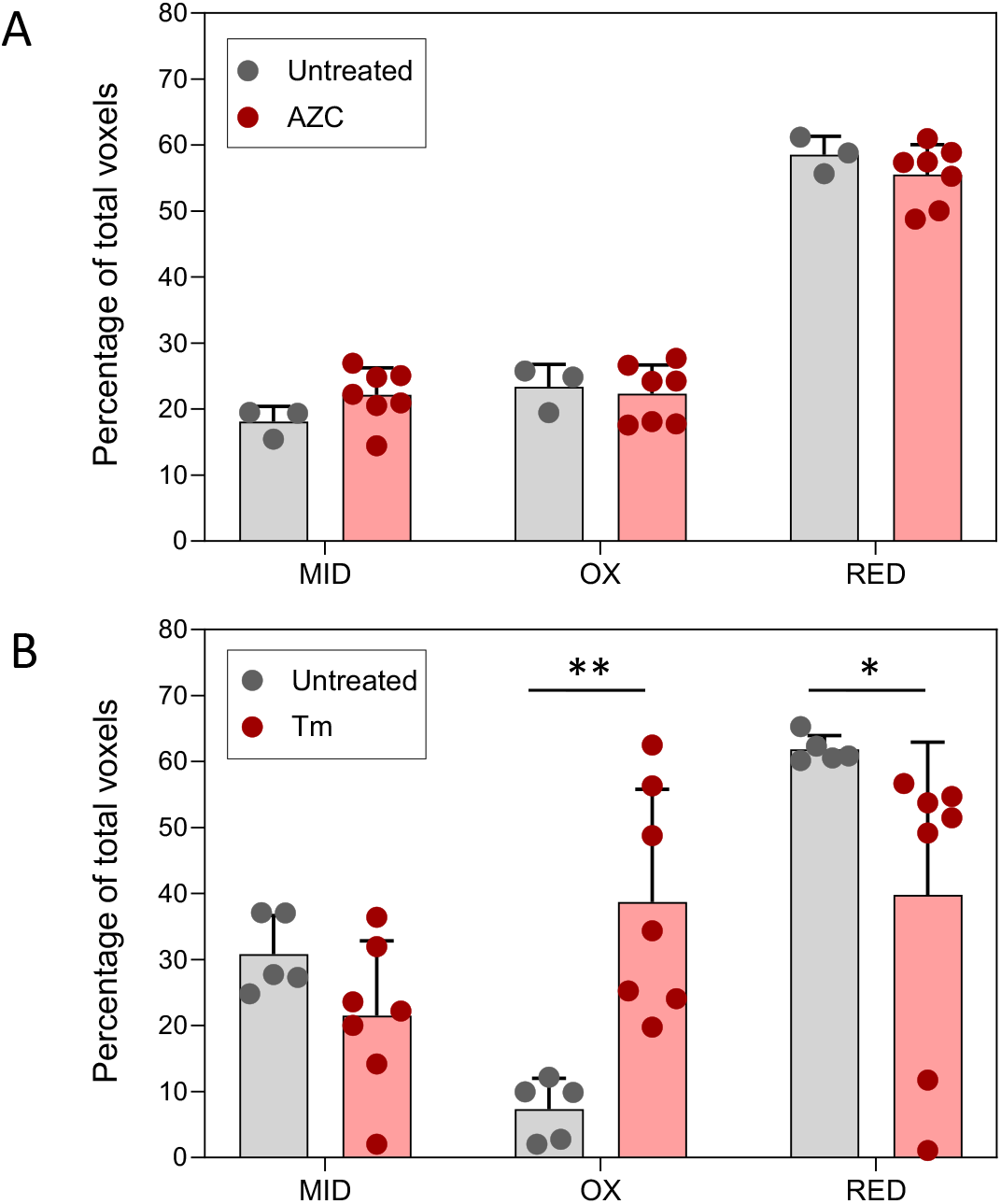
Redox state of ER is not perturbed upon proteotoxic challenges. Embryos were treated with 1 µg/ml Tm and 5 mM AZC at ~60 hpf and imaging was done after 12 hrs of treatment. Briefly, embryos were anaesthetized, mounted in lateral position imaged sequentially, i.e., collection of emission signal first upon excitation at 488 nm followed by excitation at 405 nm (confocal microscope, 10X objective). Analysis was done to calculate Ratio-iE values and percentage of voxels in each mid (intermediate), ox (oxidized), and red (reduced) bins was calculated after fitting the distributions to a 3-peak gaussian function as shown in Figure 7(A). Graph shows percentage of voxels in each bin plotted for AZC (A) and Tm (B) treated embryos with untreated embryos imaged on the same day. Each solid dot represents data from one embryo. Unpaired two tailed Student’s t test with unequal variance was performed to test significance, ** and * p-values are < 0.01 and <0.05 respectively. (n=3 and 5 for untreated in A and B respectively while n=7 for both treatments; error bars show standard deviation) (hpf-hours post fertilization).

### Limitations of the study

While we base our conclusions on our findings, there are certain limitations to our study that need to be considered for future work in this area. We have assumed that the ER-targeting signal and the ER-retention signal are equally efficient in different cell types. There is currently no such report in zebrafish or any other vertebrate systems. Since Bip (GRP78/Hspa5) is a canonical ER chaperone in all tissues, we assumed the same to be true for zebrafish. In support of this assumption, we found that Hspa5 showed ER localization patterns in all the different cells we observed in our embryonic primary culture.

## Discussion

Sub-cellular milieu plays a very important role in facilitating or obstructing the catalytic reactions and biological processes that are executed in these compartments. Accordingly, each sub-cellular compartment maintains distinct pH, ionic concentrations and redox potential appropriate to the needs of the cell. Endoplasmic reticulum is the site of protein folding, disulphide bond formation, protein secretion, calcium storage, and lipid synthesis (Schwarz & Blower, 2016). This necessitates specific conditions in the ER to facilitate these functions. For correct protein folding and disulphide formation, ER maintains an oxidizing environment. In contrast, cytosol is generally considered to be reducing. Previous studies, specifically in *C. elegans* (ER and cytosol both) and *Drosophila* (cytosol), have supported this view using in vivo redox sensors (Albrecht, et al., 2011). These and other studies in worms, *Arabidopsis thaliana*, and mice have shown that the redox potential of the cytosol is also responsive to different environmental conditions such as aging and disease (Festerling, Can, Kügler, & Müller, 2020; Kirstein, et al., 2015; Meyer et al., 2007). However, it was not clear if under physiological conditions different tissues and organs would maintain a uniform redox potential in the ER, an organelle posited to maintain only an oxidizing redox potential physiologically. The zebrafish transgenic lines with ER specific roGFP, eroGFP and ERroGFPiE, allowed us to address this intriguing question. Our studies revealed that ER redox potential varies widely across the organism. Although generally considered oxidizing, our analysis showed regions in the animal where the ER was maintained at highly reducing conditions. The cytosol also showed similar range of redox states with some parts of the embryo having oxidizing conditions.

This is the first time redox potential of the ER has been compared across an individual vertebrate organism in non-pathological conditions. The differences we observed across tissue types in ER and cytosol redox conditions suggest that our longstanding assumption that ER is maintained in an oxidized state while cytosol in a reducing state is simplistic. A large range of redox values have been reported in literature. These were determined in cell culture models and the measured values vary from −209 mV (Avezov, et al., 2013) to −231±1.87 mV (van Lith, et al., 2011) to −240 mV (in tobacco leaves; (Brach, et al., 2009)). Within a vertebrate organism we reconcile this diversity and show that cells from different regions of the embryo maintain their ER in different redox states. We believe this happens as different tissues within an organism and different cells within a tissue fulfill different biological roles. For instance, the ER in pancreatic alpha cells would primarily be folding and secreting proteins while the ER in the muscle would primarily be acting as calcium stores. This would warrant different redox milieu within these organelles in the pancreas and muscle. Our study finds very striking differences in the ER and the cytosol redox state in select regions of the brain suggesting interesting differences in cellular processes within different regions of the brain.

## Supporting information

Main tables

## Author contributions

K.C., C.S. and M.V. conceptualized and designed the work. K.C. and C.S. supervised the work. M.V. generated cyroGFP and ERroGFPiE transgenic line and performed all the zebrafish experiments. N.R.B. and K.V. generated eroGFP transgenic line. A.C. did biophysics experiments. S.U. helped in the MATLAB script writing and analysis. S.V. helped in the single cell suspension experiment for DTT and Tm treatment. Manuscript writing was done by K.C., C.S., and M.V. with inputs from all the authors. All authors read and edited the manuscript.

## Acknowledgments

We are grateful to Sir James Remington for providing us the roGFP plasmids which we used for generation of transgenic lines. We thank Dr. Sridhar Sivasubbu (CSIR-IGIB) for providing pSS550 zebrafish expression vector. This work was primarily funded by BSC0124 grant by CSIR to KC along with IGIB core funding. We thank the CIF facility of CSIR-IGIB for confocal microscope. Instrument and consumable support were also obtained from DBT-Wellcome Trust India Alliance through intermediate fellowship to K.C. MV acknowledges CSIR; NRB, SV acknowledge UGC; and AC acknowledges DBT for their fellowship.

## Competence

The authors declare no competing interests.

## *The abbreviations used in this study are

ER: endoplasmic reticulum
PDI: protein disulfide isomerase
GSH: glutathione reduced
GSSG: glutathione oxidized
dpf: days post fertilization
ss: signal sequence
a.a.: amino acids
DTT: dithiothreitol
Tm: tunicamycin
ROS: reactive oxygen species
diamide: N,N,N′,N′-tetramethylazodicarboxamide

## Materials and methods

### Zebrafish maintenance and breeding

Zebrafish were maintained and bred at standard temperature conditions of 28°C. All zebrafish experiments were performed according to the guidelines issued by the Institutional Animal Ethics Committee (IAEC) of the CSIR-Institute of Genomics and Integrative Biology, India. AB (WT), *Tg(actin:eroGFP), Tg(actin:cyroGFP)*, and *Tg(actin:ERroGFPiE)* are used in this study.

### Cloning of roGFP constructs

Conventional as well as homologous recombination-based cloning was done for generating the constructs to be used for transgenic lines. pSS550. roGFP2 and roGFPiE were amplified from pRSETB-ro2 and pQE30-ro1/iE obtained as a kind gift from Dr. James Remington lab. *Tg(actin:eroGFP)* was made from the construct made by restriction digestion based method, where 3 primers (no. 1 to 3) were used to add Hspa5 signal sequence upstream of roGFP2. Primers 4 and 5 were used to amplify the final construct and subcloned into the pSS550 vector using restriction digestion sites for NheI and ClaI. All primers used are listed in table1. For generating *Tg(actin:cyroGFP)*, roGFP2 was amplified from pRSETB-ro2 with primers 6 and 7 having 15 bp sequence homology to the vector ends obtained after restriction digestion. For *Tg(actin:ERroGFPiE)*, roGFPiE was amplified and 78 bp corresponding to Hspa5 sequence signal of 26 aa were added by sequential PCR using primers 8, 9 and 10 and 11 and 12. All primer sequences are provided in supplementary table 1. In case of ER transgenic lines, the reverse primer had KDEL sequence as well which is an ER retrieval signal. Each of these, vector and the insert, were then recombined using In-Fusion HD ® cloning kit, where 15 min reaction at 50°C was setup with enzyme mix provided in the kit. For all the 3 clonings, the reaction mix was transformed in lab made DH5α competent cells and colonies were screened the next day.

### *in vitro* transcription

Tol2 transposase was *in vitro* transcribed using pCS2 where it was first linearized by using restriction enzyme NotI. The transcript was made using SP6 RNA polymerase using mMessage mMachine kit from Invitrogen with 1 µg of template DNA. The plasmid itself had polyA at the 3’ end, so addition of polyA tailing step was not required. Quality of RNA was assessed using agarose gel electrophoresis and stored at 80°C until further use.

### Injection and generation of transgenic lines

One-cell stage embryos were collected after 15-20 min of removing the dividers. The respective roGFP construct and tol2 transposase mRNA was reconstituted in such a way that 1 nl contained 12.5 pg of each, and hence 1 nl of the volume was injected in one-cell stage embryos of WT background. Water was changed in the evening and GFP positive embryos were screened at 2 or 3 dpf stage using ZEISS AxioScope A1 microscope (with Axiocam HRc®). At least 100 GFP positive embryos were put for growing into adults, for each line. After 3 months, progeny from three independent founder lines (obtained by backcrossing with the WT line) were put for growing. Embryos obtained from at least F2 stage or more were used for experiments. Before each experiment, 1 dpf embryos were screened for GFP fluorescence using ZEISS AxioScope A1 microscope and positive embryos were processed.

### Zebrafish single cell suspension and primary culture

GFP positive (obtained from the transgenic lines) and WT embryos were first dechorionated manually and transferred to 1.5 µl microcentrifuge tubes (MCTs). 50 and 100 embryos were taken for imaging and western experiments respectively. After dechorionation, deyolking was done by adding 100 µl of lab made ringer’s solution (constitution as per zfin) for 5 min with intermittent pipetting with 200 µl tip. 1 ml of 1X trypsin-EDTA solution (GIBCO), pre warmed at 29°C, was added and kept at 29°C dry bath. 27 µl of 100 mg/ml collagenase type IV (GIBCO) was also added to it. Pipetting up and down was done using 1 ml tip for disruption of embryos into cells after every 5 min for a total incubation time of 20 min. Reaction was stopped using 200 µl FBS, kept for 2 min and the suspension was spun at 400 g for 5 min. Cells were suspended in L15 media with 10% FBS after 2 washes with chilled 1X PBS. Cell suspension was seeded either in 8-chambered coverglasses or 6-well cell culture plates (precoated with 0.2% gelatin) for imaging or western respectively. The primary culture contains all cells from embryos which could attach to the substratum. Media was changed the next day and culture was processed for further experiments.

### Localization using organelle trackers

First of all, media was removed and 10 µg/ml of Hoechst solution, made in 1X HBSS (Hank’s Balanced Saline Solution; GIBCO), was added to the primary culture in chambered slides. After 15 min, organelle trackers, working solutions made in 1X HBSS were added for 30 min. 10 µM ER-tracker, 1 µM of each Mito and LysoTracker was used. Two washes were given with 1X HBSS and the L15 media was added again. Imaging was done using 63X oil objective, 4 zoom in Leica SP8 confocal microscope.

### Protein extraction and western blotting

Embryos were harvested by removing water from the petri plates and transferred to 1.5 ml MCTs. Approximately 39 embryos were taken for western experiments. 120 µl of NP40 buffer, with PICs (Protease and Phosphatase inhibitor cocktails), was added to the embryos followed by homogenization using tissue homogenizer (name and company). Spin was given at 16,000 g for 15 min and supernatant was collected. For protein extraction from primary culture cells, cells were scraped off in PBS and collected in 1.5 ml MCTs. NP40 buffer with PICs was added and the cell suspension was vortexed for 30 min at 4°C. Spin was given at 16,000 g for 15 min and supernatant was taken. Protein concentration was estimated using BCA kit (name and company). Samples were prepared for SDS page in laemelli buffer with 30 μg of protein per sample per gel. 10-15% gels were made using premix Acrylamide:Bisacrylamide (Liqui-gel, 29:1) solution according to the need. Wet transfer of proteins was done on 0.2 μ nitrocellulose membrane (Biorad) at 70 V for 3 hrs. Probing was done with α-GFP (1:10,000; rabbit, Abcam), α-GRP78 (1:2000; rabbit, Protein tech), α-GRP94 (1:1,000; goat, enzo), α-Actin (1:2000; mouse, Sigma), α-CoxIV (1:1000, rabbit, Cell Signaling Technology), and α-SOD2 (1:1000, rabbit, abcam). Blots were developed using Crescendo solutions (name and company) in Syngene gel imager (name and company).

### ER isolation

Approximately 200 embryos were harvested for ER isolation and the protocol suggested in Nature protocols, 2013 (Prudent et al 2013; PMID not available) was followed. In brief, embryos were homogenized by tissue homogenizer in 200 μl of MB buffer. 800 μl of MB buffer was added and the mixture was spun at 1,000 g for 10 min to remove debris. The supernatant thus obtained was spun again to get nuclear fraction. This supernatant was spun again at 10,000 g for 10 min to obtain mitochondrial fraction as pellet. Spin was given again at 10,000 g for 10 min and the final supernatant was subjected to 100,000 g for 1 hr. The pellet thus obtained was considered as ER fraction and supernatant as ER supernatant. The fractions were quantified for protein concentrations and samples were made for SDS PAGE in laemelli buffer with 30 μg of protein per gel.

### Bacterial expression and purification of roGFP2 and roGFPiE

Genes for roGFP2 and roGFPiE cloned in bacterial expression plasmids pRSETB-roGFP2 (6xHIS, *Amp*^*+*^) and pQE30-roGFP1-iE (6xHIS, *Amp*^*+*^) respectively. The plasmids were transformed into *E. coli* BL21 (DE3) and single colonies were inoculated. The primary inoculums were used to inoculate 1L LB with Ampicillin (100 µg/ml) bulk cultures. Protein expression was induced with 0.7 mM IPTG upon reaching OD of 0.6. After induction, cultures were incubated for 18 hrs at 30°C, 200 rpm shaking speed. The cells were harvested and resuspended in a pre-chilled base buffer (1X PBS + 5% glycerol) containing 2 mM PMSF. Cell lysis was done using sonication and the lysates were centrifuged at 17,000 *g* for 1 h at 4°C. The supernatants thus obtained were loaded onto separate gravity-assisted Ni-NTA agarose affinity chromatography columns pre-equilibrated with chilled base buffer. The columns were washed with 10 column volumes of base buffer containing 5 mM imidazole. Proteins were eluted with 2 column volumes of base buffer containing 500 mM imidazole. The imidazole was removed by buffer exchange with base buffer at 4°C in Amicon Ultra-15 filter unit NMWL 3kDa (Millipore).

### Redox titration of roGFP2 and roGFP1-iE

Based on previous protocols, roGFP2 and roGFPiE were titrated in GFP redox buffer (75 mM HEPES + 125 mM KCl + 1 mM EDTA, pH 7.4) containing 0-10 mM DTT (DL-dithiothreitol; as a reducing agent) and 10-0 mM diamide (as an oxidizing agent) in a reciprocal manner with steps of 1 mM initially for standardizations. Then, 10 concentrations of reducing:oxidizing agents were taken between 4 to 6 mM. The titration was set up in a black opaque 96 well plate (compatible with fluorescence measurement). Final volume of redox reaction was kept at 100 µl and final concentration of GFP at 10 µM. The mixture was incubated in dark for 1 hr at room temperature. Finally, the fluorescence intensity of respective wells was measured in TECAN Infinite M200 Pro ELISA plate reader (operated by TECAN I Control V3.3.10.0 software). The excitation wavelength was set at 405 nm and 480 nm with emission wavelength kept constant at 520 nm. The data thus generated were processed and plotted in Microsoft Excel.

### DCFDA assay for ROS estimation

ROS estimation was done in 96 well plate using the cell-permeant 2’,7’-dichlorodihydrofluorescein diacetate (H_2_DCFDA) (thermo) dye and each well had 3 embryos. 12 such wells were used for one replicate of each line, so one replicate represents data from 3 embryos. Embryos in well with 10 mM H_2_O_2_ was used as a positive control. 200 μl of water was added to each well and a before reading was taken. Then excess water was removed and 200 μl of 2.5 mM DCFDA was added in each well, incubated at 37°C for 2 hrs and readings were taken in TECAN Infinite M200 Pro ELISA plate reader at 488 nm.

### Drug treatments

Tunicamycin (Sigma) was used at 6 and 10 μg/ml for 8 hrs in primary cells according to the need for imaging as well as western experiments. 0.5 mM DTT (Sigma) treatment for 8 hrs was given to embryos for western while 10 mM DTT was given, and cells were immediately imaged for redox readings within 1 min.

### Redox imaging

Embryos were molded first in low melting agarose and then redox imaging was done using leica SP8 confocal microscope with 10X objective; while 63X objective, zoom 4 was used for cells. First low melting agarose (invitrogen) was heated to boil and kept at 36°C in a dry bath for equilibration. Embryos were anaesthetized using 0.004 % Tricaine (Sigma) for 1 min or till it stopped moving, put in agarose for equilibration for a few seconds and immediately put as drop on rectangular coverslip. Embryos were molded in lateral position or according to the need using ZEISS light microscope.

Sequential scan setting for imaging was used in a confocal microscope for redox imaging. In brief, there were four settings at which embryos were imaged called channels. For embryos, channel 0-excitation (ex): 488 nm, emission (em): 505-540 nm, channel 1: bright field, channel 2: ex: 405 nm, em: 440-480 nm, and channel 3: ex: 405 nm, em: 505-540 nm. For cells the same settings were used except emission collected for excitation at 488 nm and 405 nm in channel 0 and 3 was from 505-530 nm. The obtained data was processed for image analysis. Channels were changed between frames for imaging. Here channel 2 readings were procured for subtraction of background fluorescence.

### Image analysis

Images were extracted from .lif (format of leica microscope) files in.tiff format as grey scale images. These were processed in MATLAB using in-house generated script. The pseudo code of MATLBA script is as shown in steps below:

1. Three frames of images read (image1=505/488, image2=505/405, image3 = 450/405 (emission nm/excitation nm)).
2. The intensity of a pixel is calculated as an average of surrounding pixels (1 pixel width) in the x y plane for image1 and image2.
3. All pixels where the fluorescence in image1 < CUTOFF, is replaced by zero in image1 and image2.
4. All pixels where the fluorescence in image3 > BACKGROUND_CUTOFF is replaced by zero in image1 and image2.
5. The ratio of image1/image2 is saved as the ratio file containing the ratio information for each of the pixels.
6. The pixels are colored according to the criteria set in the different conditions. The Ratio-2 images thus obtained were viewed in 3D viewer in ImageJ.

## Table Legend

**Table 1. Properties of the different peaks of Ratio-iE (shown as percentage of voxels) and redox potential as obtained from ERroGFPiE embryos.**

**Table 2: Properties of Ratio-iE and redox potential values as obtained from ERroGFPiE embryos when treated with AZC and Tm.**

## Notes

https://doi.org/10.7910/DVN/PRCFJM

### Competing Interest Statement

The authors have declared no competing interest.

### Summary of Updates

This version 2 of the manuscript has the following updated features: 1. Graphical abstract 2. Signal sequence analysis using SignalP 6.0. 3. Redox imaging data from a large number of embryos for ERroGFPiE line, n=16. 4. Redox imaging data upon Tunicamycin and AZC treatment.

