## Supplementary material for "Live visualization of endoplasmic reticulum redox potential in zebrafish embryos reveals region-specific heterogeneity": Main tables

Table 1: Properties of the different peaks of Ratio-iE (shown as percentage of voxels) and redox potential as obtained from ERroGFPiE embryos

| | | Percentage of voxels in each redox bin | | | REDOX POTENTIAL |
|---|---|---|---|---|---|
| DAY | EMBRYO TAG | MID | OX | RED | MIDDLE PEAK (mV) |
| DAY1 | embryo 1 | 19.15 | 39.30 | 41.55 | -230.50 |
| | embryo 2 | 13.95 | 39.88 | 46.17 | -231.96 |
| | embryo 3 | 21.10 | 33.97 | 44.93 | -232.15 |
| | embryo 4 | 18.11 | 34.22 | 47.66 | -229.99 |
| DAY2 | embryo 1 | 24.82 | 9.88 | 65.30 | -237.03 |
| | embryo 2 | 37.08 | 2.78 | 60.14 | -234.09 |
| | embryo 3 | 27.31 | 12.16 | 60.54 | -234.21 |
| | embryo 4 | 27.74 | 9.95 | 62.31 | -234.86 |
| | embryo 5 | 37.10 | 2.02 | 60.87 | -234.34 |
| DAY3 | embryo 1 | 19.35 | 19.44 | 61.21 | -237.58 |
| | embryo 2 | 15.43 | 25.78 | 58.79 | -236.32 |
| | embryo 3 | 19.49 | 24.85 | 55.66 | -235.47 |
| DAY4 | embryo 1 | 28.05 | 18.54 | 53.41 | -235.65 |
| | embryo 2# | 61.74 | 34.57 | 3.69 | -228.45 |
| | embryo 3 | 23.10 | 25.08 | 51.82 | -236.47 |
| | embryo 4 | 26.84 | 28.54 | 44.63 | -236.22 |
| AVERAGE | | 26.27 | 22.56 | 51.17 | -234.08 |
| SD | | 11.23 | 12.03 | 14.22 | 2.63 |

### This embryo exhibits an outlier distribution.

**COLUMN LEGEND**

DAY: Experiments done on different days are labelled as separate sets

EMBRYO TAG: each embryo done on a day is serially labelled.

OX: Percentage of voxels exhibiting oxidizing redox potential

RED: Percentage of voxels exhibiting reducing redox potential

MID: Percentage of voxels exhibiting intermediate redox potential

MIDDLE PEAK (mV): Calculated redox potential of the intermediate peak (MID) in milli volts.

SD: standard deviation

TABLE 2: Properties of Ratio-iE and redox potential values as obtained from ERroGFPiE embryos when treated with AZC and Tm.

| | | Percentage of voxels in each redox bin | | | REDOX POTENTIAL |
|---|---|---|---|---|---|
| TREATMENT | EMBRYO TAG | MID | OX | RED | MIDDLE PEAK (mV) |
| AZC | embryo 1 | 24.82 | 17.75 | 57.43 | -236.33 |
| | embryo 2 | 26.95 | 24.25 | 48.80 | -237.17 |
| | embryo 3 | 20.92 | 18.12 | 60.96 | -236.94 |
| | embryo 4 | 20.53 | 24.19 | 55.29 | -235.06 |
| | embryo 5 | 25.08 | 17.57 | 57.35 | -235.17 |
| | embryo 6 | 14.46 | 26.64 | 58.90 | -236.68 |
| | embryo 7 | 22.26 | 27.68 | 50.06 | -235.23 |
| | AVERAGE | 22.14 | 22.31 | 55.54 | -236.08 |
| | SD | 3.82 | 4.06 | 4.19 | 0.84 |
| Tunicamycin | embryo 1 | 31.96 | 56.30 | 11.75 | -237.14 |
| | embryo 2 | 2.02 | 48.80 | 49.18 | -267.89 |
| | embryo 3 | 23.58 | 19.77 | 56.65 | -234.67 |
| | embryo 4 | 36.42 | 62.52 | 1.07 | -264.06 |
| | embryo 5 | 20.03 | 25.26 | 54.71 | -233.41 |
| | embryo 6 | 14.20 | 34.34 | 51.46 | -234.35 |
| | embryo 7 | 22.19 | 24.07 | 53.74 | -235.12 |
| | AVERAGE | 21.49 | 38.72 | 39.79 | -243.81 |
| | SD | 10.50 | 15.82 | 21.42 | 14.10 |

**COLUMN LEGEND**

TREATMENT: Experiments labelled with the chemicals used to perturb proteostasis. Either AZC (azetidine-2-carboxylic acid) at 5 mM or Tunicamycin at 1 μg/ml were used to treat the embryos for 12 hours.

EMBRYO TAG: each embryo done on a day is serially labelled.

OX: Percentage of voxels exhibiting oxidizing redox potential

RED: Percentage of voxels exhibiting reducing redox potential

MID: Percentage of voxels exhibiting intermediate redox potential

MIDDLE PEAK (mV): Calculated redox potential of the intermediate peak (MID) in milli volts.

SD: standard deviation
