## Supplementary file for "Live visualization of endoplasmic reticulum redox potential in zebrafish embryos reveals region-specific heterogeneity"

### Supplemental Information

### Live visualization of cytosolic and endoplasmic reticulum redox potential in zebrafish embryos reveals region-specific differences

Monika Verma, Niraj Rajesh Bhatt, Aseem Chaphalkar, Kriti Verma, Shreyansh Umale, Shweta Verma, Chetana Sachidanandan, and Kausik Chakraborty

#### Inventory of supplemental items

1. Figure S1 - independent
2. Figure S2 - independent
3. Figure S3 - relates to main Figure 1
4. Figure S4 - relates to main Figure 1
5. Figure S5 - independent
6. Figure S6 - independent
7. Figure S7 - relates to main Figure 2
8. Figure S8 - relates to main Figure 4
9. Figure S9 - relates to main Figure 5 and 6
10. Figure S10 - relates to main Figure 7
11. Table S1- independent

Supplementary figure 1

A

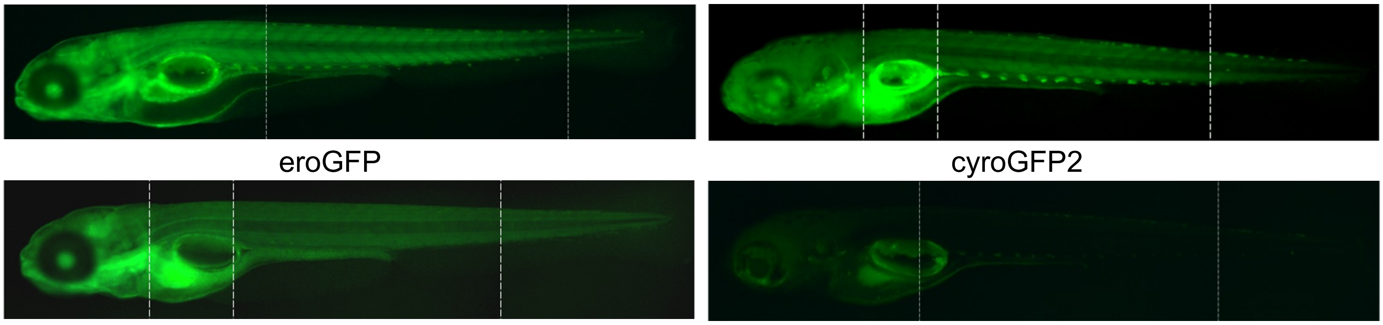


**Supplementary figure 1: roGFP is expressed ubiquitously in all transgenic lines.** (A) 4 dpf embryos were anaesthetized and mounted in lateral position in 2.5% methyl cellulose. show. Representative fluorescence images of the three transgenic lines and wild type is shown (epifluorescence microscope, 5X objective, dpf- days post fertilization).

Supplementary figure 2

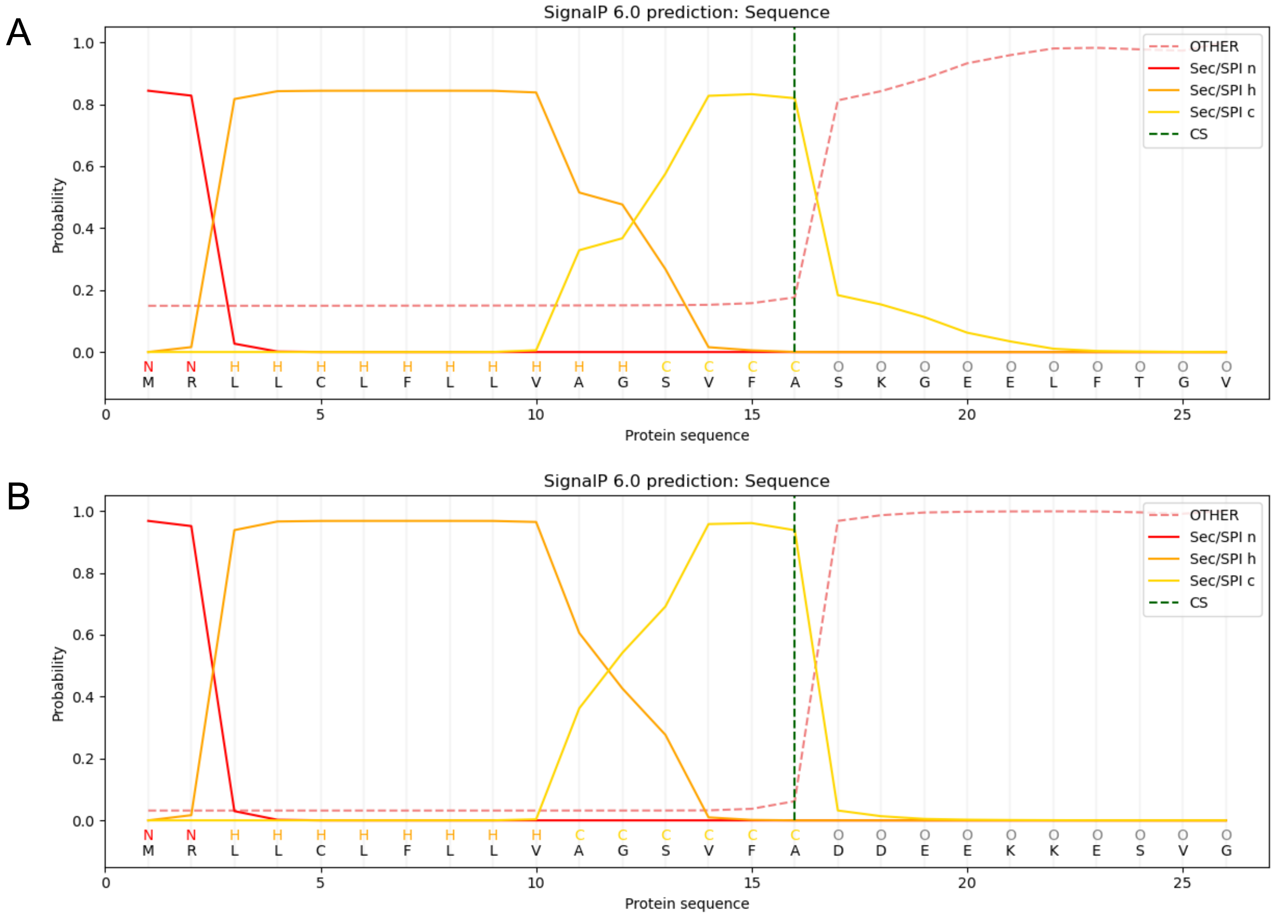


**Supplementary figure 2: Signal peptide cleavage prediction revealed proper cleavage of both the signal sequences used.** We used free online software, SignalP 6.0, to predict signal peptide cleavage site for the signal sequences we used for roGFP transgenic lines. (A) First 16 a.a. of Hspa5 (GRP78) followed by first 10 a.a. (after removing methionine) of roGFP2 were used for the prediction. (B) First 26 a.a. of Hspa5 (GRP78) were used for prediction. The black dotted line is the predicted residue after which cleavage would be done. (a.a.- amino acid, CS- cleavage site, Sec/SPI- transported by Sec translocon and cleaved by Signal Peptidase I, n h and c are N-terminal, hydrophobic and C-terminal regions of signal peptide)

Supplementary figure 3

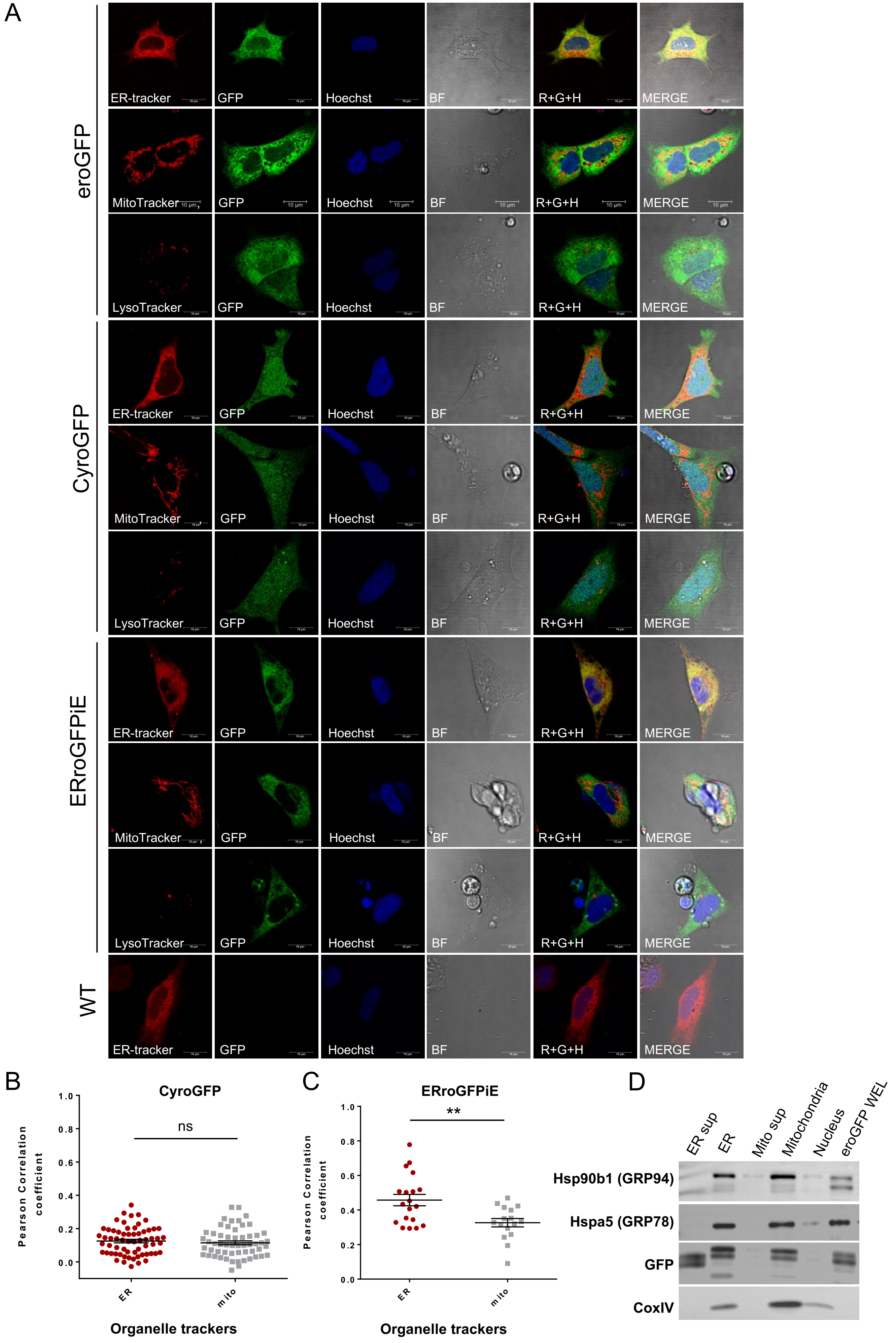

**Supplementary figure 3: roGFP probes are localized in the desired organelles in generated transgenic lines.** (A) Single cell suspension was made form 1 dpf embryos and cells were attached to substratum (8 chambered coverglasses) in L15 medium at 28°C. Imaging was done after overnight attachment of live cells with different stainings. Fluorescence image of zebrafish primary culture cells with ER, Mito and LysoTracker staining for all the transgenic lines are shown, with wild type fish as a negative control. These panels correspond to the images of transgenic lines as labeled. The first image in each panel is ER, Mito or LysoTracker followed by the GFP channel corresponding to roGFP. Third image is hoechst staining for nucleus and next image is bright field (confocal microscope, 63X objective, 4X zoom, scale 10μm). (B) and (C) Scatter plot shows Pearson correlation coefficient of green and red channel for cyroGFP and ERroGFPiE line respectively, where each dot is one ROI. (D) The eroGFP embryos were harvested at 5 dpf and ER isolation was done by subcellular fractionation which was done using sucrose and differential centrifugation. Western blots show fractions obtained after fractionation. Hspa5 (GRP78) and Hsp90b1 (GRP94) were used as ER markers while CoxIV as a mitochondrial marker along with probing for GFP as well. Two tailed Student's t test with unequal variance was done to test significance, ** is p value <0.01. (BF- bright field, R+G+H- merge of red, green and hoechst channel, ROI- region of interest, ER- endoplasmic reticulum, mito- mitochondria, sup- supernatant, WEL- whole embryo lysate)

Supplementary figure 4

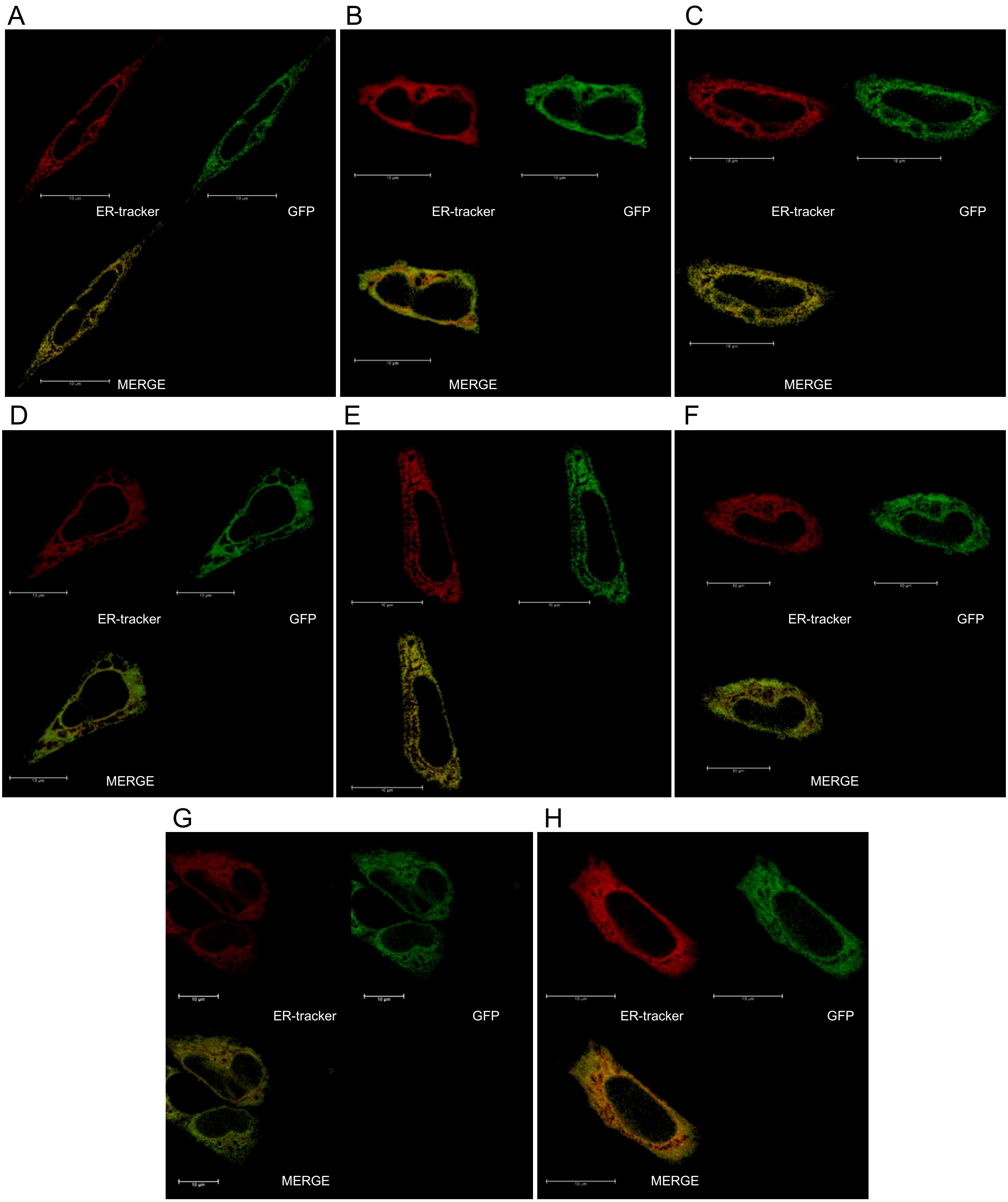


**Supplementary figure 4: Representative images for ER Tracker Red colocalization with roGFP2 in eroGFP line.** Fluorescence images of zebrafish primary culture cells with ER Tracker staining for eroGFP line are shown. The first image in upper panel is ER Tracker Red, followed by the GFP channel corresponding to roGFP2. Image in lower panel image is merge of red and green channel showing yellow pixels (confocal microscope, 63X objective, 4X zoom, scale 10μm).

Supplementary figure 5

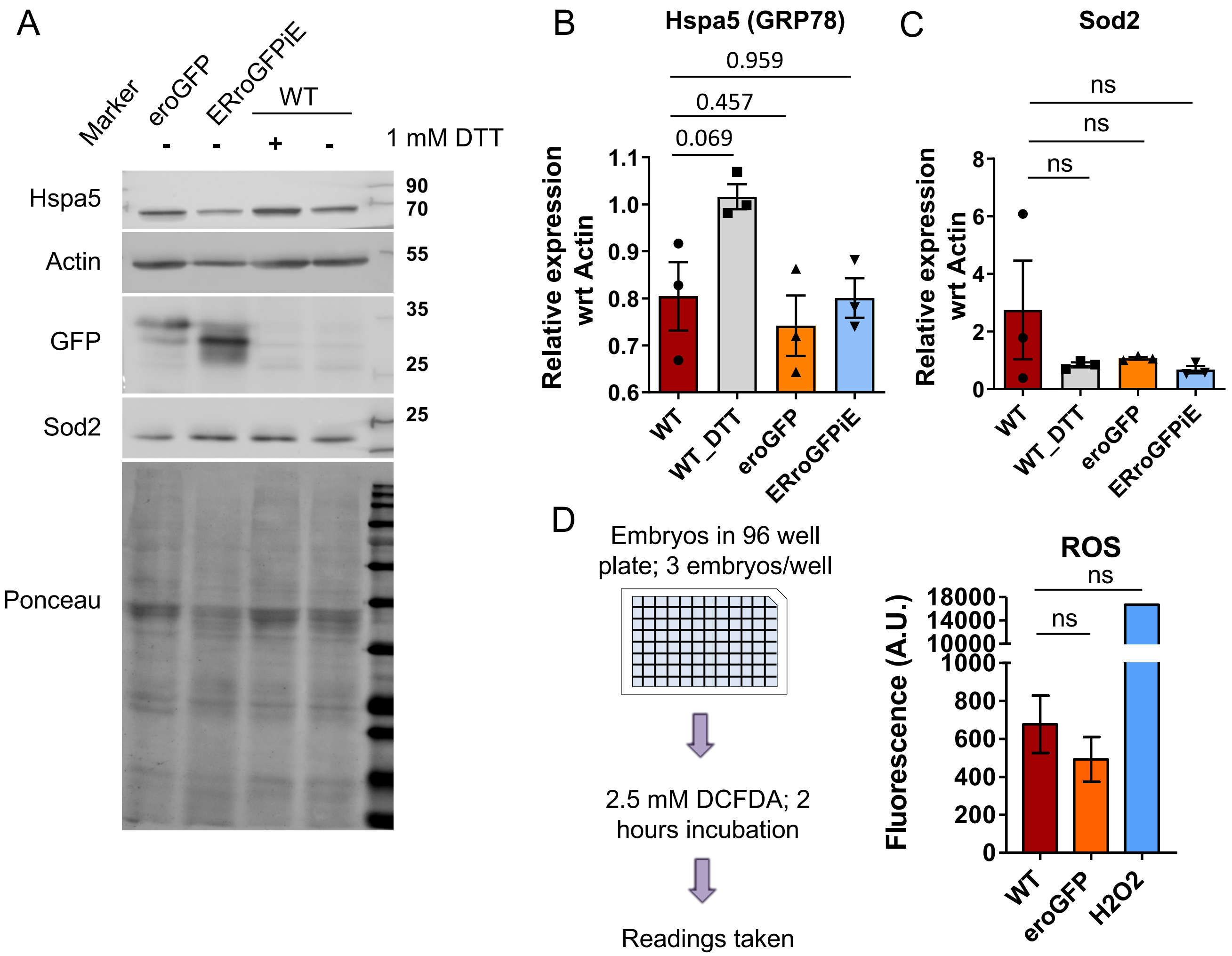


**Supplementary figure 5: roGFP transgenic lines for ER do not exhibit perturbations in homeostasis.** (A) Protein lysates from 39 embryos were loaded on a SDS gel and the transfer was done. Western blots show expression of Hspa5 (GRP78) and Sod2 in the ER transgenic lines and wild type along with 1mM DTT treated embryos (8 hrs) taken as a positive control. GFP was also probed, and actin was used as a loading control. (B) and (C) The graph shows quantification of Hspa5 and Sod2 expression w.r.t. actin respectively which was done using ImageJ. Paired Student's t-Test with two tails was performed using Microsoft excel. The bars show mean while error bars show SEM. (D) 3 live embryos at 5 dpf were put in each well of a 96 well plate. Water was removed to an extent that embryos have some water to survive and DCFDA dye was added. Incubation was done for 2 hrs and readings were taken. Wells with embryos incubated in $H_2O_2$ was used as a positive control. 12 wells were used for one replicate, so each replicate had 36 embryos. Unpaired Student's t-Test wad done with Welch correction using Graph pad Prism. Bars represents mean from 3 replicates while error bars are SEM (WT- wild type, w.r.t- with respect to, SEM-standard error around mean)

Supplementary figure 6

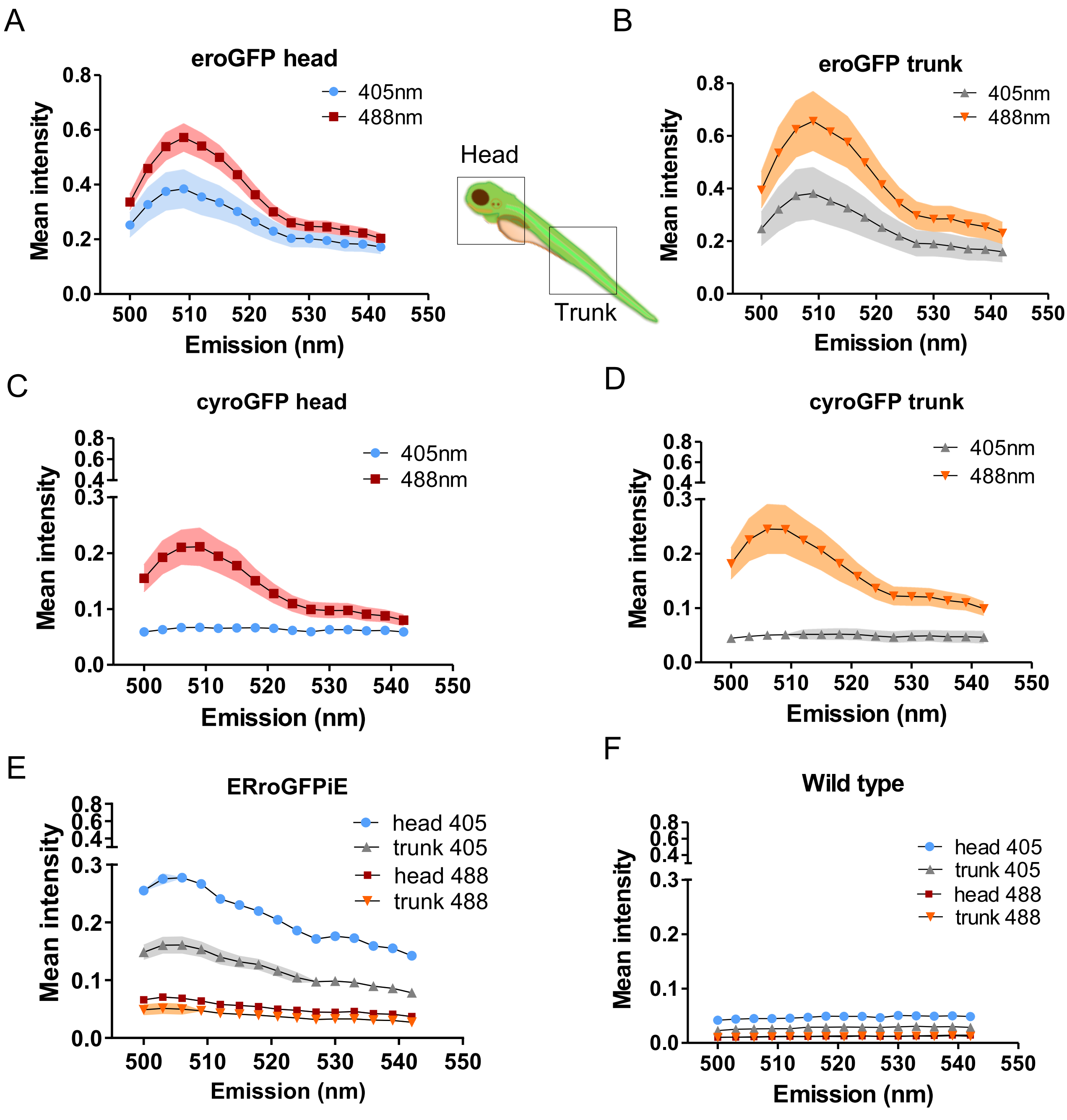


**Supplementary figure 6: roGFP transgenic lines show redox sensitive properties *in vivo.*** Embryos were excited at first 405 nm and emission scan was taken from 500 nm to 540 nm, then excited at 488 nm and emission was collected in the same range. These graphs show emission scan for live embryos at 4 dpf where head and trunk regions are marked in the cartoon representation of an embryo. (A) and (B) Graphs show emission peaks when excited at 405 and 488 nm for head and trunk regions of eroGFP line respectively. (C), and (D) Graphs show emission peaks at 405 and 488 nm for head and trunk regions of cyroGFP line respectively (E) and (F) Peaks at 405 and 488 nm for ERroGFPiE and WT lines are shown respectively, settings were kept same for all. Each line represents mean value from 4 replicates and shaded region indicates SEM. (SEM- standard error around mean)

Supplementary figure 7

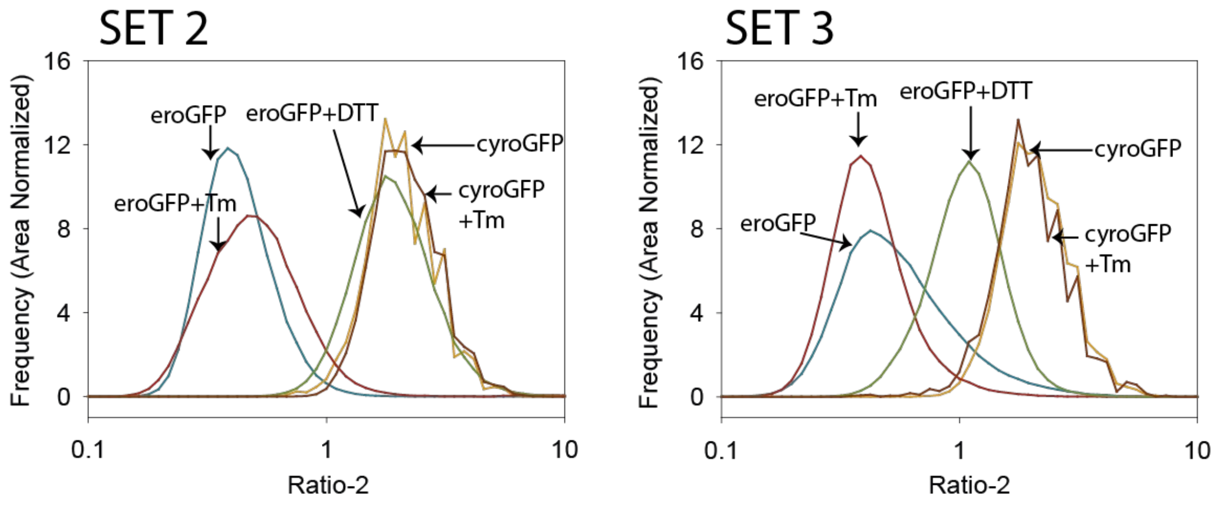


**Supplementary figure 7: Ratio-2 distribution of primary cell culture from 1 dpf zebrafish embryos.** 1 dpf zebrafish embryos were processed to get single cell suspension followed by growing in L15 medium in chambered coverglasses. Cells kept overnight were treated with 6µg/ml Tm (8 hrs) and 10 mM DTT (1-2 min) and immediately imaged sequentially to obtain emission signals (from 505-540 nm) upon excitation at 488 nm and 405 nm. Ratio-2 values obtained after image analysis were plotted separately for each condition in both Sets. Each Set and each line represents data from one replicate and mean Ratio-2 for each treatment respectively. N is variable in each condition and contains value from at least 6 fields.

Supplementary figure 8

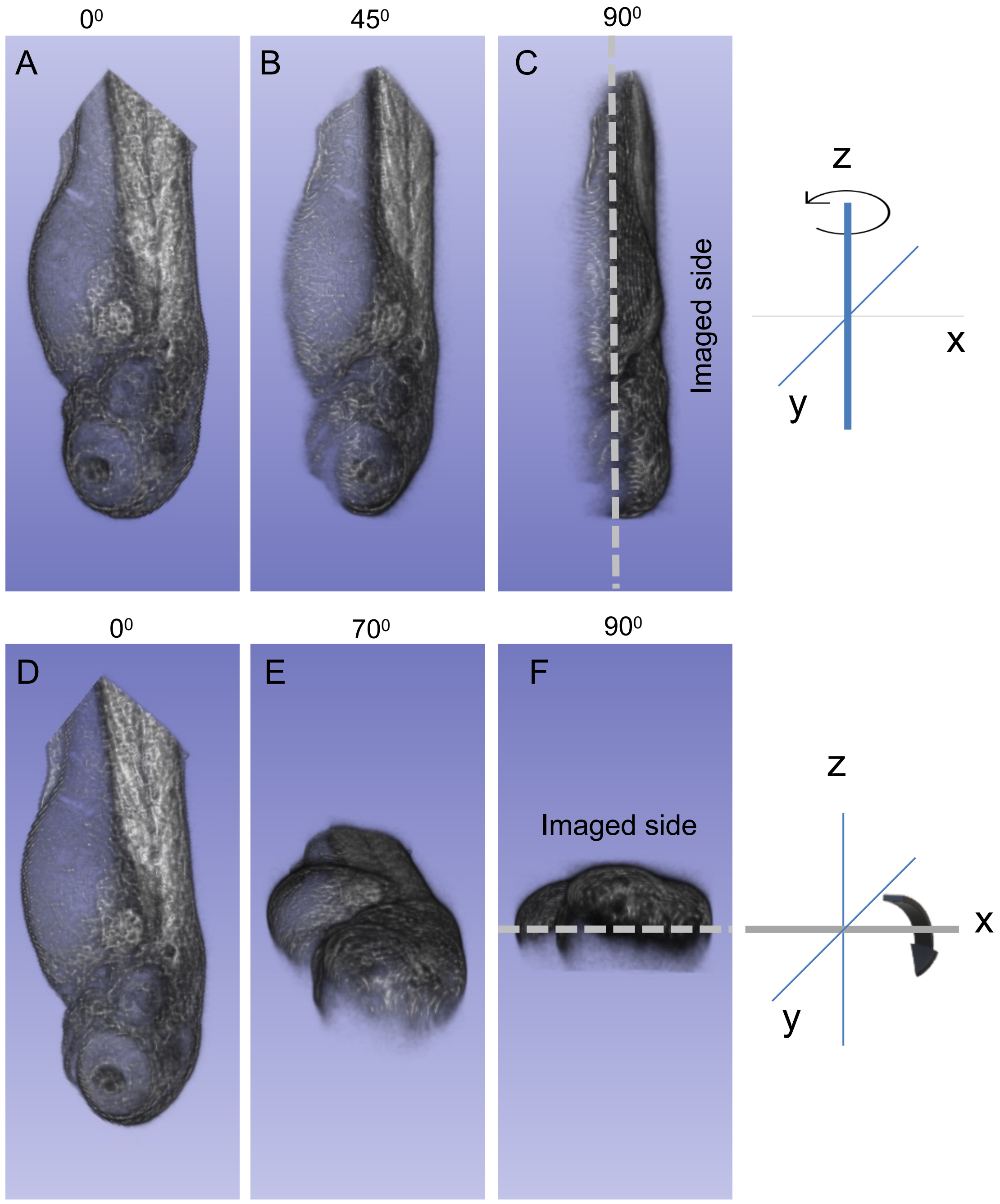


**Supplementary figure 8: Confocal imaging enables to procure signals from either half of the embryo in 3D.** 3D representation of a 3 dpf zebrafish embryo was reconstructed from z-stack images of GFP fluorescence. A single embryo is shown for illustrating the orientations used in all the figures. (A) and (D) is the $0^0$ angle of image showing the way embryo was imaged. (B) and (C) represents the embryo at $45^0$ and $90^0$ respectively when turned left to right as indicated in the arrow. (E) and (F) represents the embryo at $70^0$ and $90^0$ respectively when turned up from down (shown in arrow). (Confocal microscope, 10X objective)

Supplementary figure 9

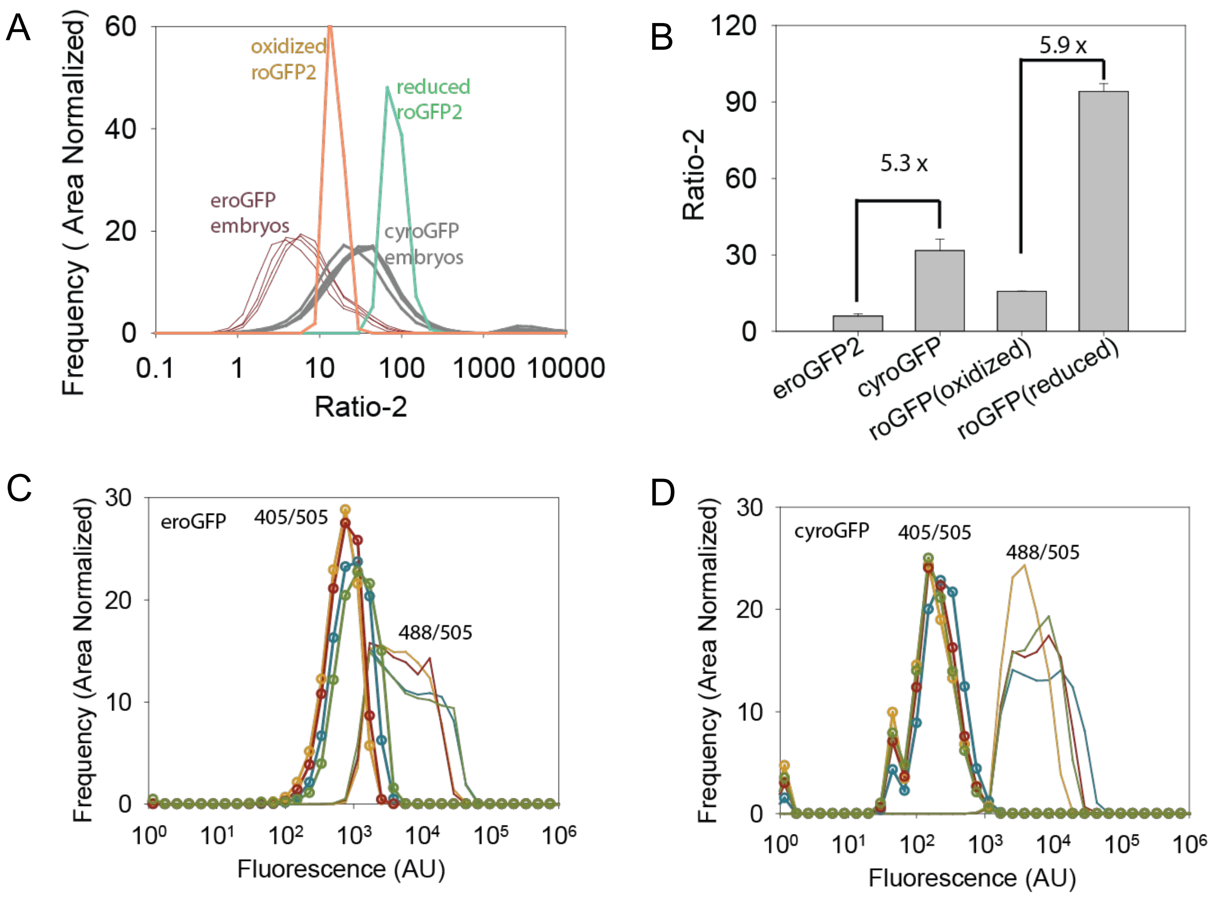


**Supplementary figure 9: Ratio-2 distribution of eroGFP and cyroGFP fish lines showed shift in the peak as compared to the oxidized and reduced roGFP2 solution.** 3 dpf embryos were anaesthetized, mounted on soft agar, and imaged for obtaining emission signals (505-540 nm) when excited at 488 nm and 405 nm sequentially. 5 μM roGFP2 solution was treated with 10 mM DTT and 10 mM diamide to get fully oxidized and reduced species. These solutions were also imaged using similar settings. Ratio-2 values were calculated using MATLAB. (A) The frequency of each value was area normalized using the total number of pixels and graph was plotted (x axis in log10 scale). (B) Bar graph showing mean ratio-2 for the embryos as well as roGFP2 solution (oxidized and reduced) and error bars represent standard deviation. (C) and (D) Fluorescence intensities obtained with excitation at 405 nm and emission at 505 nm (405/505) and excitation at 488 nm and emission at 505 nm (488/505) were calculated for embryos. Frequency (area normalized) was plotted against fluorescence intensity for 405/505 and 488/505 for eroGFP (left panel) and cyroGFP (right panel) embryos. Each line represents one embryo in (A), (B), and (D). (number of embryos, n= 4; confocal microscope, 10X objective).

Supplementary figure 10

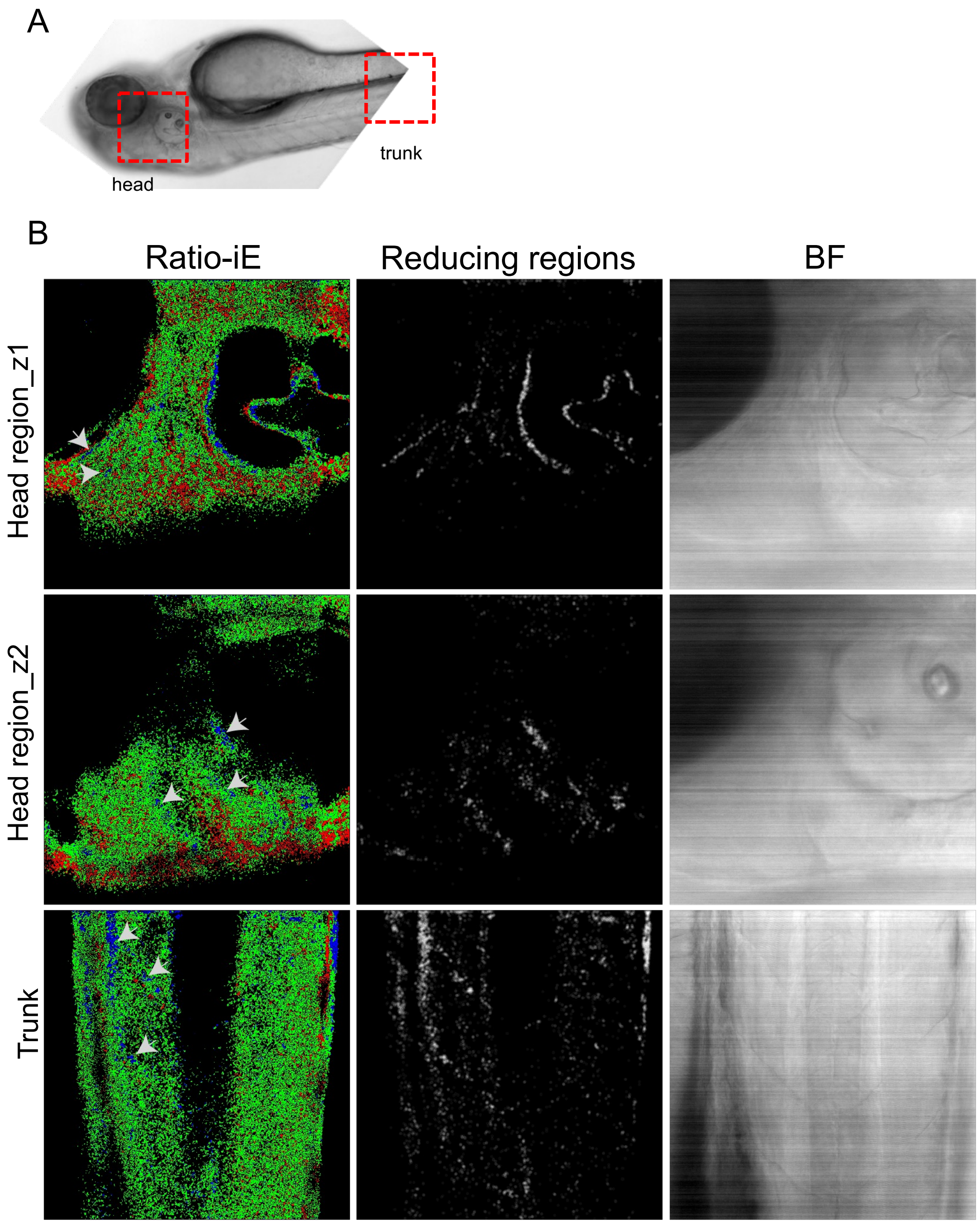


**Supplementary figure 10: Variations in redox ratios of ER are more apparent with higher magnification images of ERroGFPiE embryos.** 3 dpf live ERroGFPiE embryos were anaesthetized, mounted in soft agar, imaged, analyzed. Ratio-iE values were calculated using MATLAB and color coding was done as described by ERroGFPiE embryos. (A) A 10X BF image of an embryo showing head and trunk regions (red squares) imaged at higher magnification of 40X. (B) All three panels show representative image from one embryo taken at different regions. Left, middle, and right panel show Ratio-iE, only reducing pixels, and BF images respectively. Upper panel and middle panel shows the Ratio-iE in the eye region (black region on top left) images taken at different z depths inside the embryo. Some distinct reducing regions are marked with white arrows. Lower panel shows the Ratio-iE images taken at the trunk regions showing patches of reduced regions (white arrows) (BF- bright field, confocal microscope, 40X objective).

Supplementary table 1. Primers used for the generation of transgenic lines

| S. no. | Primer name | Primer sequence (5'-3') | Description |
|---|---|---|---|
| 1 | zf.er.erogfp.infr1 | Tttttgctggtggccggcagcgtgtttgccat ggtgagcaagggcgaggagctgttcacc | fusing hspa5 ER SS with roGFP |
| 2 | zf.er.erogfp.fr | Agtcagaattcatgcggttgctttgcctgttttt gctggtggccggcagcgtgtttgcc | Amplification of zebrafish eroGFP forward primer |
| 3 | er.erogfp.rv | Actaagcggccgcttacaattcgtcgtgctt gtacaattcgtccataccg | Amplification of roGFP, reverse primer |
| 4 | zf.er.erogfp.NotI.fr | Agtcagcggccgcatgcggttgctttgcctg tttttgctggtggccggcagcgtgtttgcc | Amplification of zebrafish eroGFP, final forward primer |
| 5 | zf.er.erogfp.ClaI.rv | Actaaatcgatttacaattcgtcgtgcttgta caattcgtccataccg | Amplification of zebrafish eroGFP, final reverse primer |
| 6 | CyroGFP2_pSS550 _for_ZF | ccatctagagcggccatgagtaaaggaga agaacttttca | Amplification of cyroGFP2, forward primer |
| 7 | CyroGFP2_pSS550 _rev_ZF | tctggatcatcatcgttatttgtatagttcatcc atgcca | Amplification of cyroGFP2, reverse primer |
| 8 | ERroGFPiE_pSS550 _in_for1_ZF | Gacgataagaaggagagtgttgggagta aaggagaagaacttttcac | fusing hspa5 ER SS with roGFP1/iE, inner forward primer |
| 9 | ERroGFPiE_pSS550 _for2_ZF | Gtggccggcagcgtgtttgccgaagagga cgataagaaggagagtgttgg | fusing hspa5 ER SS with roGFP1/iE, 2nd forward primer |
| 10 | ERroGFPiE_pSS550 _for3_ZF | Atgcggttgctttgcctgtttttgctggtggcc ggcagcgtgtttgccga | fusing hspa5 ER SS with roGFP1/iE, 3rd forward primer |
| 11 | ERroGFPiE_pSS550 _out_for4_ZF | ccatctagagcggccatgcggttgctttgcc tg | Amplification of zebrafish roGFP2/iE, final forward primer |
| 12 | ERroGFPiE_new_R ev_pSS_ZF | Tctggatcatcatcgctacagctcgtcctttt gtatagttcatccatgc | Amplification of zebrafish roGFP2/iE, final reverse primer |
